# TOMM40 regulates hepatocellular and plasma lipid metabolism via an LXR-dependent pathway

**DOI:** 10.1101/2024.06.27.600910

**Authors:** Neil V Yang, Justin Y Chao, Kelly A Garton, Tommy Tran, Sarah King, Joseph Orr, Jacob H Oei, Alexandra Crawford, Misun Kang, Reena Zalpuri, Danielle Jorgens, Pranav Konchadi, John S Chorba, Elizabeth Theusch, Ronald M Krauss

## Abstract

The gene encoding TOMM40 (Transporter of Outer Mitochondrial Membrane 40) is adjacent to that encoding APOE, which has a central role in lipid and lipoprotein metabolism. Human genetic variants near *APOE* and *TOMM40* are strongly associated with plasma lipid levels, but a specific role for TOMM40 in lipid metabolism has not been established. Investigating this, we show that suppression of *TOMM40* in human hepatoma cells upregulates expression of *APOE* and *LDLR* in part via activation of LXRB (NR1H2) by oxysterols, with consequent increased uptake of VLDL and LDL. This is in part due to disruption of mitochondria-endoplasmic reticulum contact sites, with resulting accrual of reactive oxygen species and non-enzymatically derived oxysterols. With *TOMM40* knockdown, cellular triglyceride and lipid droplet content are increased, effects attributable in part to receptor-mediated VLDL uptake, since lipid staining is significantly reduced by concomitant suppression of either *LDLR* or *APOE*. In contrast, cellular cholesterol content is reduced due to LXRB-mediated upregulation of the ABCA1 transporter as well as increased production and secretion of oxysterol-derived cholic acid. Consistent with the findings in hepatoma cells, *in vivo* knockdown of *TOMM40* in mice results in significant reductions of plasma triglyceride and cholesterol concentrations, reduced hepatic cholesterol and increased triglyceride content, and accumulation of lipid droplets leading to development of steatosis. These findings demonstrate a role for TOMM40 in regulating hepatic lipid and plasma lipoprotein levels and identify mechanisms linking mitochondrial function with lipid metabolism.

**GRAPHICAL ABSTRACT:** 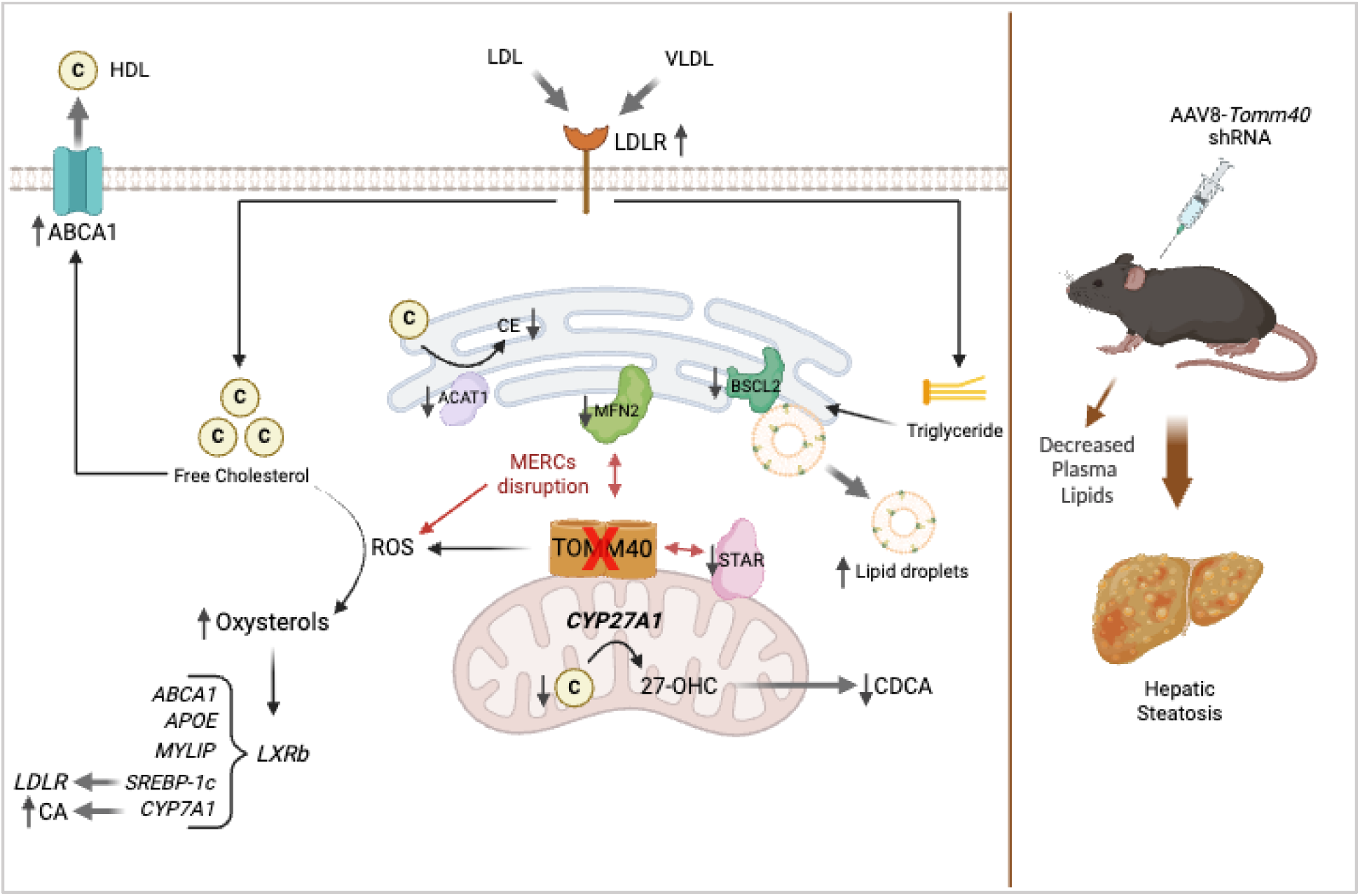

## INTRODUCTION

TOMM40 (translocase of outer mitochondrial membrane 40) is the main channel-forming subunit of the translocase of the outer mitochondrial membrane (TOM) complex (**Fig 1A**). This complex is required for the transport of precursor proteins into the mitochondria to maintain mitochondrial function^1^. Furthermore, TOMM40 has been shown to interact with key regulators of mitochondrial function such as BAP31^2^, affect calcium signalling via its interaction with VDAC channels (voltage dependent anion channels)^3–5^, and have effects at mitochondria-ER contact sites (MERCs) that impact mitochondrial cholesterol transport via steroidogenic acute regulatory protein (STAR aka STARD1)^6,7^. In mouse Leydig cells, the absence of TOMM40 causes STAR to lose its ability as a cholesterol transporter, depleting mitochondrial cholesterol content^8^.

**Figure 1.**
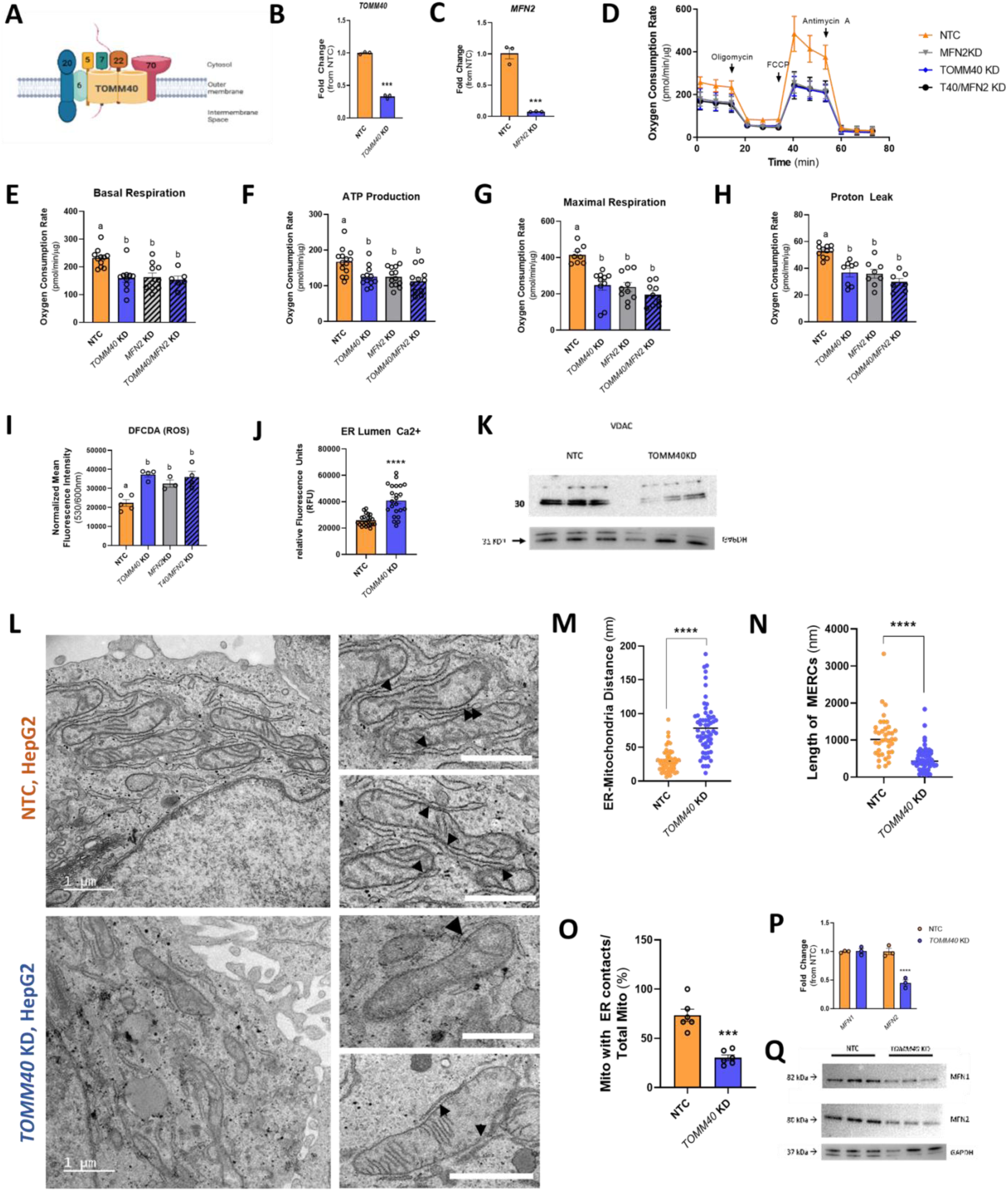
TOMM40 is essential for maintaining mitochondrial function and MERCs in hepatocytes. (A) Schematic diagram of the mammalian TOM complex, consisting of 7 subunits, located in the outer mitochondrial membrane. (B) Confirmation of *TOMM40* KD in HepG2 human hepatoma cells by ∼75%, measured by qPCR. (*n=3* biological replicates) (C) Confirmation of *MFN2* KD in HepG2 cells by ∼90%, measured by qPCR. (*n=3* biological replicates) (D-H) (D) Oxygen consumption rates of HepG2 cells transfected with *TOMM40*, *MFN2*, and/or NTC siRNAs were quantified using the Seahorse 96e Extracellular Flux Analyzer. With the addition of oligomycin, FCCP, and Antimycin A + Rotenone, basal respiration (E), ATP production (F), maximal respiration (G), and proton leak (H) were quantified. (*n=10-12* biological replicates) (I) Cellular ROS in *TOMM40* and *MFN2* KD vs. NTC HepG2 cells was quantified by DFCDA fluorescence probe. (*n=4-6* biological replicates) (J) ER Lumen Ca2+ levels were measured by Mag-Fluo-4 AM fluorescence probe. (*n=12* biological replicates) (K) Representative western blot of VDAC protein expression in HepG2 cells compared to GAPDH control. (L) TEM micrographs of NTC and TOMM40 KD in HepG2 cells. Arrowheads indicate MERCs; scale bars, 1μm. (M-O) Analysis of MERCs using ImageJ software: (M) ER-mitochondria distance (nm), (N) length of MERCs (nm), (O) percentage of mitochondria with ER contacts out of total mitochondria per cell. (*n= 12-24* cells/group) (P) mRNA transcript levels of *MFN1* and *MFN2* in HepG2 cells quantified by qPCR. (*n=3* biological replicates) (Q) Representative western blot of MFN1 and MFN2 protein expression in HepG2 cells compared to GAPDH control. For all: *p<0.05, **p<0.01, ***p<0.005, ****p<0.001 vs. NTC by one-way ANOVA, with post-hoc Student’s t-test to identify differences between groups. p<0.05 for *a* vs. *b* by two-way ANOVA, with Sidak’s multiple comparisons test. Data are represented as mean ± SEM.

The *TOMM40* gene shares the same locus on chromosome 19q as *APOE,* a gene with a key role in cholesterol and lipoprotein metabolism^9,10^. Importantly, APOE acts as a key ligand for low-density lipoprotein receptor (LDLR) to transport lipoproteins into the liver^11^. Genome-wide association studies have revealed multiple genetic variants in a haplotype block encompassing *TOMM40* and *APOE* that are associated with plasma lipid levels^12,13^. However, a specific role for TOMM40 in regulation of hepatic lipid and plasma lipoprotein metabolism has not previously been determined, and the present study was aimed at testing this possibility.

## RESULTS

### Loss of hepatic *TOMM40* disrupts mitochondrial function and mitochondria-ER contact sites (MERCs)

Consistent with prior studies in HeLa cells and *C. Elegans*^14,15^, knockdown (KD) of *TOMM40* in human hepatoma HepG2 cells (**Fig 1B**) resulted in reduction of basal and maximal respiration and ATP production as well as proton leak (**Fig 1D-H**) compared to non-targeting control (NTC). The impact of *TOMM40* KD on mitochondrial function was also manifested by an increase in cellular reactive oxygen species (ROS) (**Fig 1I**), consistent with effects reported in epithelial ovarian cancer cells^16^.

Notably, there was an increase of calcium ions (Ca^2+^) in the ER lumen with *TOMM40* KD, suggesting a block in the transfer of Ca^2+^ ions from ER to mitochondria (**Fig 1J**), likely due to a reduction in expression of VDACs (voltage-dependent anion channels) that import Ca^2+^ into mitochondria^17^ (**Fig 1K**). This finding, in conjunction with previous evidence that TOMM40 interacts directly with BAP31 at mitochondria-ER contact sites (MERCs) in U2OS osteosarcoma cells^2^, led us to test the effect of *TOMM40* on MERCs in HepG2 cells. Transmission electron microscopy (TEM; **Fig 1L**) revealed disruption of MERCs through a significant increase in ER-mitochondria distance (**Fig 1M**), decrease in length of MERCs (**Fig 1N**), and percentage of mitochondria with ER contacts (**Fig 1O**). Consistent with these effects, *TOMM40 KD* resulted in decreased protein expression of mitofusin 1 and 2 (MFN1 and MFN2) which play roles in tethering ER to mitochondria and maintaining MERCs^18^, as well as reduced *MFN2* gene expression (**Fig 1P-Q**). Many studies have shown suppression of *MFN2* to result in disruption of MERCs as assessed by electron microscopy in various human cell lines^19–21^. Thus, we further demonstrated that disruption of MERCs by KD of *MFN2*, with a knock-down efficiency of ∼90% (**Fig 1C**), reduced mitochondrial oxidative phosphorylation (**Fig 1E-H**) and increased cellular ROS to a similar extent as seen with *TOMM40* KD, and that with their combined KD, no further increase occurred (**Fig 1I**). Taken together, these results show that TOMM40 is a key regulator of mitochondrial function and MERCs and suggest that MERC disruption mediates the reduction in mitochondrial function and the increase in cellular ROS by *TOMM40* KD.

### *TOMM40* KD promotes production of oxysterols and upregulation of LXR gene targets

In addition to the increase in ROS induced by *TOMM40* KD in HepG2 cells, we observed increased levels of several cholesterol oxidation products. While oxysterols generated from ROS, such as 7-ketocholesterol, were elevated (**Fig 2A**), we also saw an increase in those produced by enzymatic reactions, namely 25- and 24(S)-hydroxycholesterol (OHC; **Fig 2B-C**). In addition, gene expression of *CYP3A4*, which encodes the rate limiting enzyme for generating 4β-hydroxycholesterol, also increased in the *TOMM40* KD cells (**Fig 2D**). Oxysterols are potent activators of the LXRA and LXRB transcription factors^22^, and therefore we next tested whether *TOMM40* KD in HepG2 cells resulted in increased expression of the LXRs and their downstream gene targets. We observed that *TOMM40* KD induced a significant increase in gene expression of *LXRB* (aka *NR1H2*), but not *LXRA* (aka *NR1H3*) (**Fig 2E**), suggesting *TOMM40* KD and its downstream ligands favor the LXRB isoform. Consistent with both LXR activation and increased *LXRB* expression, we found that transcripts of five LXR-regulated genes involved in lipid metabolism (*APOE, ABCA1, CYP7A1*, *F2BF1c,* and *MYLIP*) were significantly upregulated by *TOMM40* KD (**Fig 2F-J)**. Moreover, addition of exogenous 25-OHC to NTC-treated cells to induce LXR activation yielded effects consistent with those of *TOMM40* KD, implying an oxysterol-dependent mechanism (**Fig 2K-L**). Interestingly, double KD of *TOMM40* together with either *LXRA* or *LXRB* resulted in a reciprocal increase of the other isoform, thus maintaining relatively constant total *LXR* gene expression (**Fig S1**). Such a compensatory mechanism between the two isoforms has been reported previously^23^. We therefore tested the effects of *TOMM40* KD on expression of the five LXR target genes in conjunction with inhibition of both *LXRA* and *LXRB* by GSK2033, an LXR antagonist with high binding affinity for LXRA and LXRB, and found that all were reduced to levels similar to those seen with NTC (**Fig 2F-J**). Together, these results indicate that suppression of *TOMM40* upregulates expression of LXR target genes both by oxysterol activation of LXR and upregulation of *LXRB* to a greater extent than *LXRA*.

**Figure 2.**
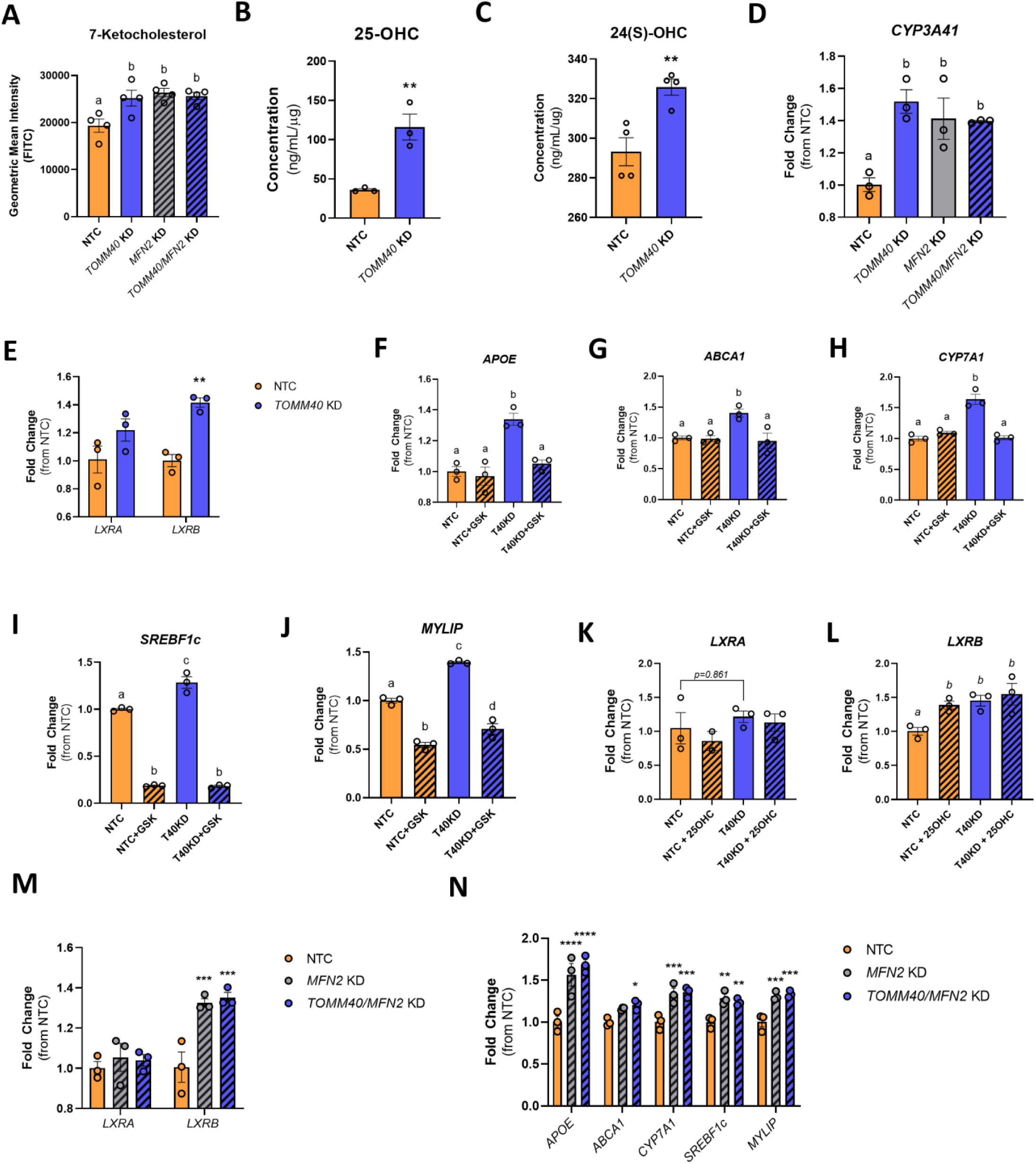
*TOMM40* KD upregulates LXRB and downstream gene targets by promoting oxysterol production. (A) 7-ketocholesterol levels in HepG2 cells transfected with NTC vs. *TOMM40, MFN2, or TOMM40/MFN2* siRNAs by flow cytometry. (*n=4* biological replicates) (B-C) Analysis of enzymatic-derived 25-OHC levels and 24(S)-OHC in *TOMM40* KD vs. NTC HepG2 cells by ELISAs. (*n=3* biological replicates) (D) mRNA transcript levels of *CYP3A4*, responsible for the synthesis of 4β-OHC, in HepG2 cells, quantified by qPCR. (*n=3* biological replicates) (E-J) (E) mRNA transcript levels of *LXRA* and *LXRB* and their downstream targets: *APOE* (F), *ABCA1* (G), *CYP7A1* (H*), SREBF1c* (I), and *MYLIP* (J), in HepG2 cells quantified by qPCR. (K-L) mRNA transcript levels of *LXRA* (K) and *LXRB* (L) in HepG2 cells transfected with NTC vs. TOMM40 siRNAs after the addition of 10 μM GSK2033 (LXR antagonist). (*n=3* biological replicates) (M-N) (M) mRNA transcript levels of *LXRA* and *LXRB* and their downstream targets (N) in NTC, *TOMM40* KD, *MFN2* KD, and *TOMM40/MFN2* KD HepG2 cells, quantified by qPCR. (*n=3* biological replicates) For all: *p<0.05, **p<0.01, ***p<0.005, ****p<0.001 vs. NTC by one-way ANOVA, with post-hoc Student’s t-test to identify differences between groups. p<0.05 for *a* vs. *b* vs. *c* vs. *d* by two-way ANOVA, with Sidak’s multiple comparisons test. Data are represented as mean ± SEM.

Moreover, we found that *MFN2* KD also upregulated expression of *LXRB* (**Fig 2M**) and LXR target genes (**Fig 2N**) similar to *TOMM40* KD, and increased cellular concentration of 7-ketocholesterol (**Fig 2A**) and *CYP3A4* expression (**Fig 2D**), but not enzymatic-derived oxysterols (**Fig S2**). This suggests that ROS-derived 7-ketocholesterol is a key ligand driving LXR activity via disruption of MERCs in both *MFN2* KD and *TOMM40* KD HepG2 cells.

### *TOMM40* KD increases LDLR gene expression and receptor-mediated LDL hepatic uptake

Notably, we found that in addition to the known LXR transcriptional targets, *LDLR* gene expression in HepG2 cells was also increased by *TOMM40* KD (**Fig 3A**), as were cellular LDLR protein content (**Fig 3B**), and cell surface LDLR (**Fig 3C**). Furthermore, the increase of *LDLR* was reversed by addition of GSK2033, indicating a role for LXRs in *LDLR* regulation (**Fig 3A**). Expression of *SREBF2*, the canonical transcriptional regulator of *LDLR*, was unaffected by *TOMM40* KD, consistent with unaffected transcript levels of *HMGCR* (**Fig S3**), another downstream target of SREBF2. However, we found that *TOMM40* KD-induced upregulation of *LDLR* gene expression was suppressed by concurrent KD of *SREBF1c*, a known transcriptional target of LXR^24^ and splice variant of *SREBF1*(**Fig 3D**). Moreover, gene expression of *LDLR* (but not *HMGCR*) was suppressed by *SREBF1c* KD alone **(Fig 3D).**

**Figure 3.**
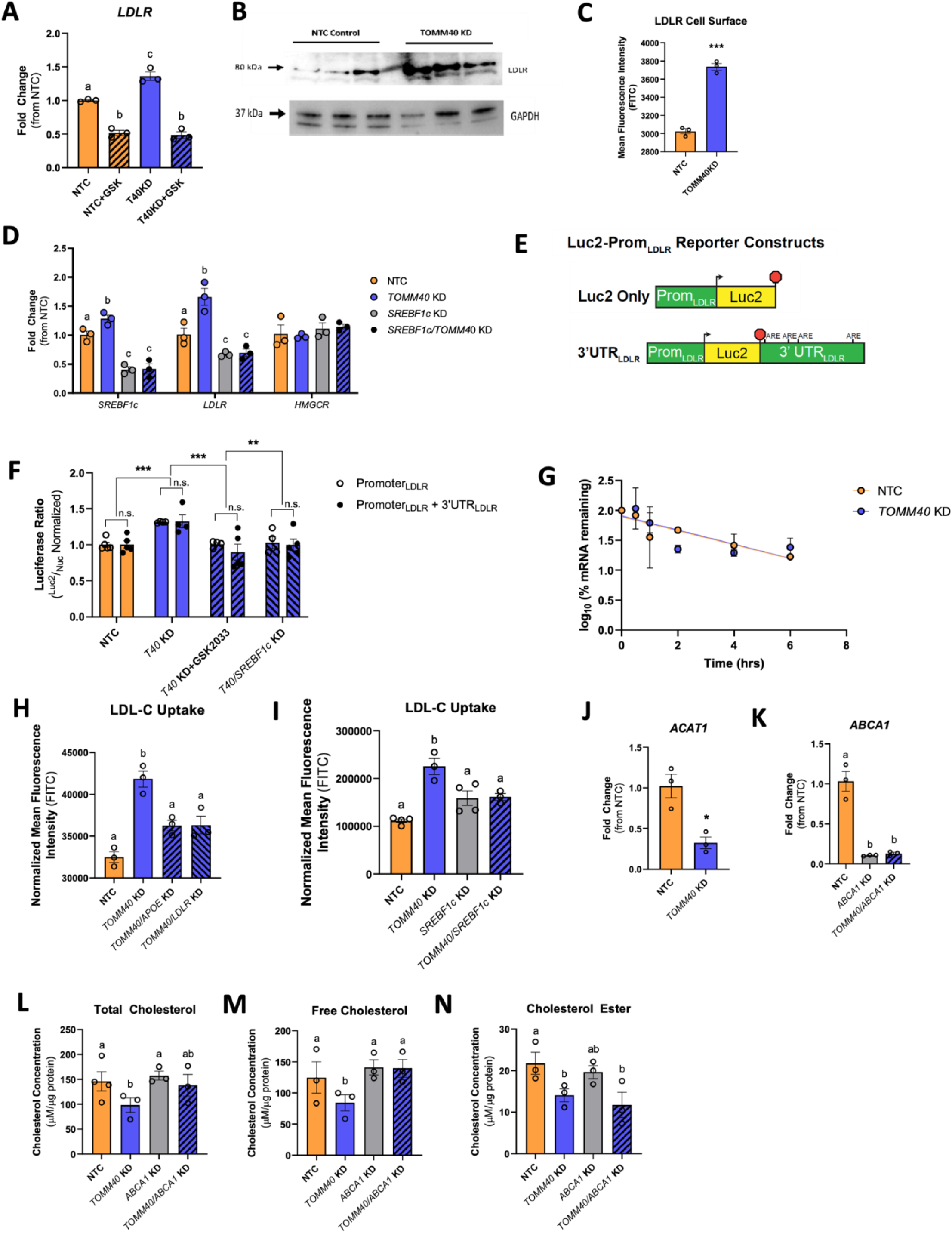
*TOMM40* KD promotes LDL uptake via an LXR-mediated pathway. (A) *LDLR* mRNA transcripts were quantified by qPCR in HepG2 cells treated with NTC vs. *TOMM40* KD, with or without GSK2033. (*n=3* biological replicates) (B) Representative western blot of LDLR protein expression in NTC vs. *TOMM40* KD HepG2 cells. (C) LDLR cell surface protein levels were stained with anti-LDLR antibody and analyzed by flow cytometry. (*n=3* biological replicates) (D) mRNA transcripts confirming KD of *SREBF1c* and changes in expression of *LDLR* and *HMGCR* in *SREBF1*, *TOMM40*, and NTC siRNAs-treated HepG2 cells. (*n=3* biological replicates) (E) Schematic diagram of Luc2-Prom_LDLR_ reporter constructs, illustrating *LDLR* promoter, start site (arrowhead), stop codon (red octagon), and 3’UTR region, containing adenylate-uridylate (AU)-rich elements (AREs) implicated in mRNA stability, of the *LDLR* gene. (F) Ratiometric luciferase outputs of HepG2 cells transfected with indicated reporters. (*n=4-5* biological replicates) n.s. = non-significant. Two-way ANOVA, **p<0.01, ***p<0.005 by Holm-Sidak test. Data are represented as mean ± SEM. (G) Relative expression of *LDLR* mRNA in NTC vs. *TOMM40* KD HepG2 cells after arrest of transcription with actinomycin D. (*n=3* biological replicates) (H-I) BODIPY-labelled LDL-C was taken up in HepG2 cells and fluorescence was quantified on a microplate fluorescence spectrophotometer. (*n=3* biological replicates) (J-K) mRNA transcripts of *ACAT1* (J) and *ABCA1* (K) expression quantified by qPCR. (L-N) Intracellular total cholesterol (L), free cholesterol (M), and cholesterol ester (N) levels quantified from HepG2 cells transfected with NTC, *TOMM40*, and *ABCA1* siRNAs, singly and in combination, using Amplex Red Cholesterol Assay. (*n=3-4* biological replicates) For all (except F): *p<0.05, **p<0.01, ***p<0.005, ****p<0.001 vs. NTC by one-way ANOVA, with post-hoc Student’s t-test to identify differences between groups. p<0.05 for *a* vs. *b* vs. *c* by two-way ANOVA, with Sidak’s multiple comparisons test. Data are represented as mean ± SEM.

To further assess the transcriptional relationship between *SREBF1c* and *LDLR* expression with *TOMM40* KD, we transiently expressed luciferase constructs containing the native *LDLR* promoter with and without the *LDLR* 3’UTR in *TOMM40* KD HepG2 cells (**Fig 3E**). While *TOMM40* KD alone increased LDLR promoter activity, this returned to normal in the presence of GSK2033 or *SREBF1c* KD (**Fig 3F**). No differences in activity were observed between constructs fused to the LDLR promoter + 3’UTR coding sequence vs. the LDLR promoter alone. Furthermore, we showed no effect of *TOMM40* KD on *LDLR* mRNA decay rate and stability after 4 hrs treatment with 1μg/mL of Actinomycin D (**Fig 3G**). Taken together, these results indicate thar the upregulation of *LDLR* transcript by *TOMM40* KD is mediated by LXR-driven expression of *SREBF1c* targeting the *LDLR* promoter region.

We next demonstrated that *TOMM40* KD increased LDL uptake in HepG2 cells (**Fig 3H**), and that this effect was blocked by KD of either *LDLR* or *APOE*, indicating that it is likely due to increased expression of these genes via LXR activation. Moreover, we showed that KD of *SREBF1c* inhibited the increase in LDL uptake by *TOMM40* KD to an extent similar to that of its reduction by *LDLR* KD, adding evidence for an LXR – SREBF1c – *LDLR* mechanism (**Fig 3I**). Unexpectedly however, *TOMM40* KD markedly decreased content of intracellular cholesterol (free and esterified) (**Fig 3L-N**). Notably, KD of *ABCA1*, which encodes a transporter responsible for cellular cholesterol efflux and is a transcriptional target of LXR, (**Fig 3K)** resulted in restoration of cellular (total and free) cholesterol content in *ABCA1/TOMM40* KD HepG2 cells (**Fig 3L-N**). Moreover, with *TOMM40* KD there was reduced gene expression of *ACAT1* (**Fig 3J**) which encodes the enzyme required for esterification of free cholesterol^25^. Therefore, through LXR activation and upregulation of LXRB, *TOMM40* KD promotes LDL uptake, but in parallel increases cholesterol efflux via ABCA1 and reduces ACAT1-mediated cholesterol esterification, resulting in reduced intracellular cholesterol levels.

### Loss of *TOMM40* expression promotes the classic bile acid synthesis pathway, while inhibiting the alternative pathway via STAR

Since oxysterols serve as precursors of bile acid synthesis, we tested whether *TOMM40* KD-induced increases in oxysterol levels resulted in increased total bile acid secretion in HepG2 cells^26^.While we found a greater amount in the media of *TOMM40* KD vs. NTC cells (**Fig 4A**), there was no change in intracellular levels (**Fig 4B**). We then measured cellular content of the two major bile acids: cholic acid (CA) and chenodeoxycholic acid (CDCA) of the classical and alternative bile synthesis pathways, respectively^27^, and found that *TOMM40* KD increased the level of CA in both cells (**Fig 4C**) and media (**Fig 4D**), whereas it decreased intracellular CDCA (**Fig 4E,F**), and this was primarily responsible for no net change in total intracellular bile acids. The effect on CA is consistent with our finding that *TOMM40* KD upregulates LXR-mediated expression of *CYP7A1*, which encodes the rate-limiting enzyme in the classic bile acid synthesis pathway^28^. These results indicate that the increase in total bile acid synthesis and secretion with *TOMM40* KD is driven at least in part by CYP7A1 of the classic bile acid synthesis pathway.

**Figure 4.**
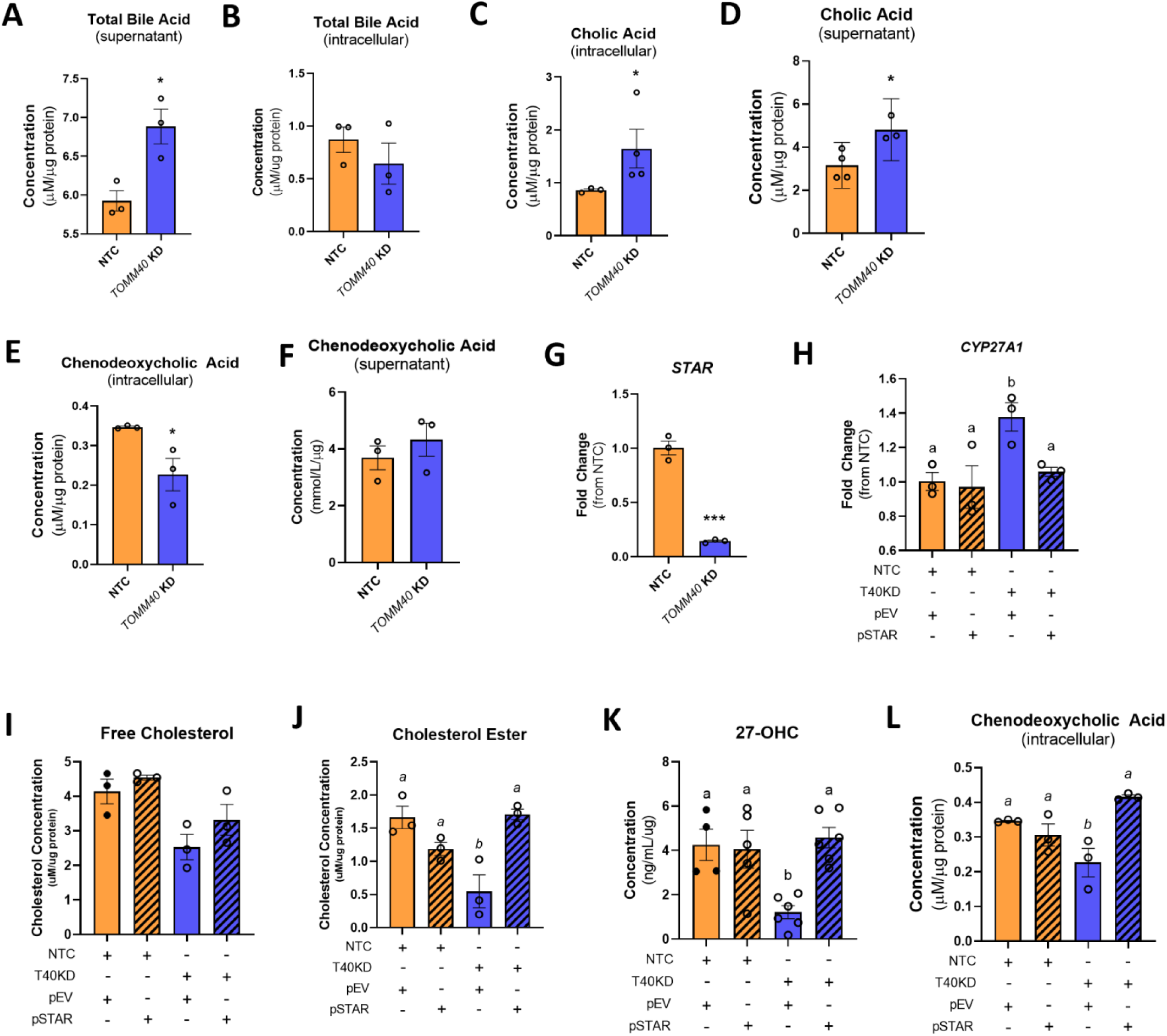
*TOMM40* KD in hepatocytes promotes classic bile acid synthesis pathway while inhibiting alternative pathway via interaction with STARD1 at MERCs. (A-B) Total bile acid levels in NTC vs. *TOMM40* KD HepG2 cells were measured in the supernatant (cell media; A) and intracellularly (B) by ELISA and normalized to protein concentration by BCA assay. (C-D) Cholic acid levels were quantified intracellularly (C) and in the supernatant (D) of HepG2 cells. (E-F) Chenodeoxycholic acid levels were measured intracellularly (E) and in the supernatant (F) in NTC vs. *TOMM40* KD HepG2 cells. (G) mRNA transcripts of *STAR* were quantified in NTC vs. *TOMM40* KD HepG2 cells by qPCR. (H) *CYP27A1* gene expression was measured in NTC vs. TOMM40 KD HepG2 cells with overexpression of an empty vector (pEV) or a *STAR*-expressing plasmid (pSTAR). (I-J) Mitochondrial free cholesterol (I) and cholesterol ester (J) levels were measured by Amplex Red Cholesterol Assay in HepG2 cells. (K) Analysis of enzymatic-derived 27-OHC levels in *TOMM40* KD vs. NTC HepG2 cells overexpressed with pEV or pSTAR by ELISA. (L) Analysis of intracellular chenodeoxycholic acid levels in NTC vs. *TOMM40* KD HepG2 cells overexpressing pEV or pSTAR. For all: *p<0.05, **p<0.01, ***p<0.005, ****p<0.001 vs. NTC by one-way ANOVA, with post-hoc Student’s t-test to identify differences between groups. p<0.05 for *a* vs. *b* by two-way ANOVA, with Sidak’s multiple comparisons test. Data are represented as mean ± SEM. (*n=3* biological replicates)

To assess the basis for reduced cellular CDCA content with *TOMM40* KD, we measured expression of *CYP27A1*, which encodes 27-cholesterol hydroxylase, the first enzyme in the alternative bile acid pathway that converts cholesterol into 27-OHC in the inner mitochondrial membrane^29^. Surprisingly, we found that its mRNA expression was increased by *TOMM40* KD (**Fig 4H**) whereas both mitochondrial cholesterol content (**Fig 4I,J**) and 27-OHC cellular levels (**Fig 4K**) were reduced, thus providing an explanation for the reduction in cellular CDCA, the downstream product of 27-OHC, despite upregulation of *CYP27A1* transcripts. In *MFN2* KD HepG2 cells, 27-OHC levels were also significantly reduced, suggesting reduction in 27-OHC and CDCA to be mediated by MERCs (**Fig S4**). Having observed depletion of mitochondrial cholesterol content, we next assessed the role of STAR (aka STARD1), a mitochondrial cholesterol transporter located at MERCs^30^, and found that its gene expression was suppressed in *TOMM40* KD HepG2 cells (**Fig 4G**). Moreover, overexpression of *STAR* rescued the reduction of mitochondrial cholesterol content by *TOMM40* KD (**Fig 4I,J**) and increased cellular levels of 27-OHC (**Fig 4K**) and CDCA (**Fig 4L**) levels similar to those seen with NTC. Furthermore, *STAR* overexpression reversed the upregulation of *CYP27A1* gene expression by *TOMM40* KD in HepG2 cells (**Fig 4H**). These findings demonstrate that the interaction between TOMM40 and STAR at MERCs plays a role in regulating mitochondrial cholesterol content and that depletion of mitochondrial cholesterol content by *TOMM40* KD suppresses the alternative bile acid synthesis pathway.

### Effects of *TOMM40* KD on hepatic triglyceride metabolism

In contrast to the depletion of cholesterol by *TOMM40* KD in HepG2 cells, we found that triglyceride content was significantly increased (**Fig 5A**). We therefore tested the effect of *TOMM40* KD on VLDL uptake and showed an increase that was prevented by *LDLR* suppression, similar to what was observed for LDL uptake (**Fig 5B**). We further assessed the effect of *TOMM40* KD on expression of other receptors involved in hepatic VLDL uptake and showed upregulation of *SDC1 and LRP1*, but not *VLDLR* (**Fig 5C**). Unlike the *LDLR* KD results, KD of *TOMM40* in combination with KD of either *SDC1* or *LRP1* did not reduce VLDL uptake (**Fig 5D**). Moreover, only the *TOMM40* and *LDLR* double KD cells showed a significant reduction in intracellular triglyceride levels (**Fig 5E**).

**Figure 5.**
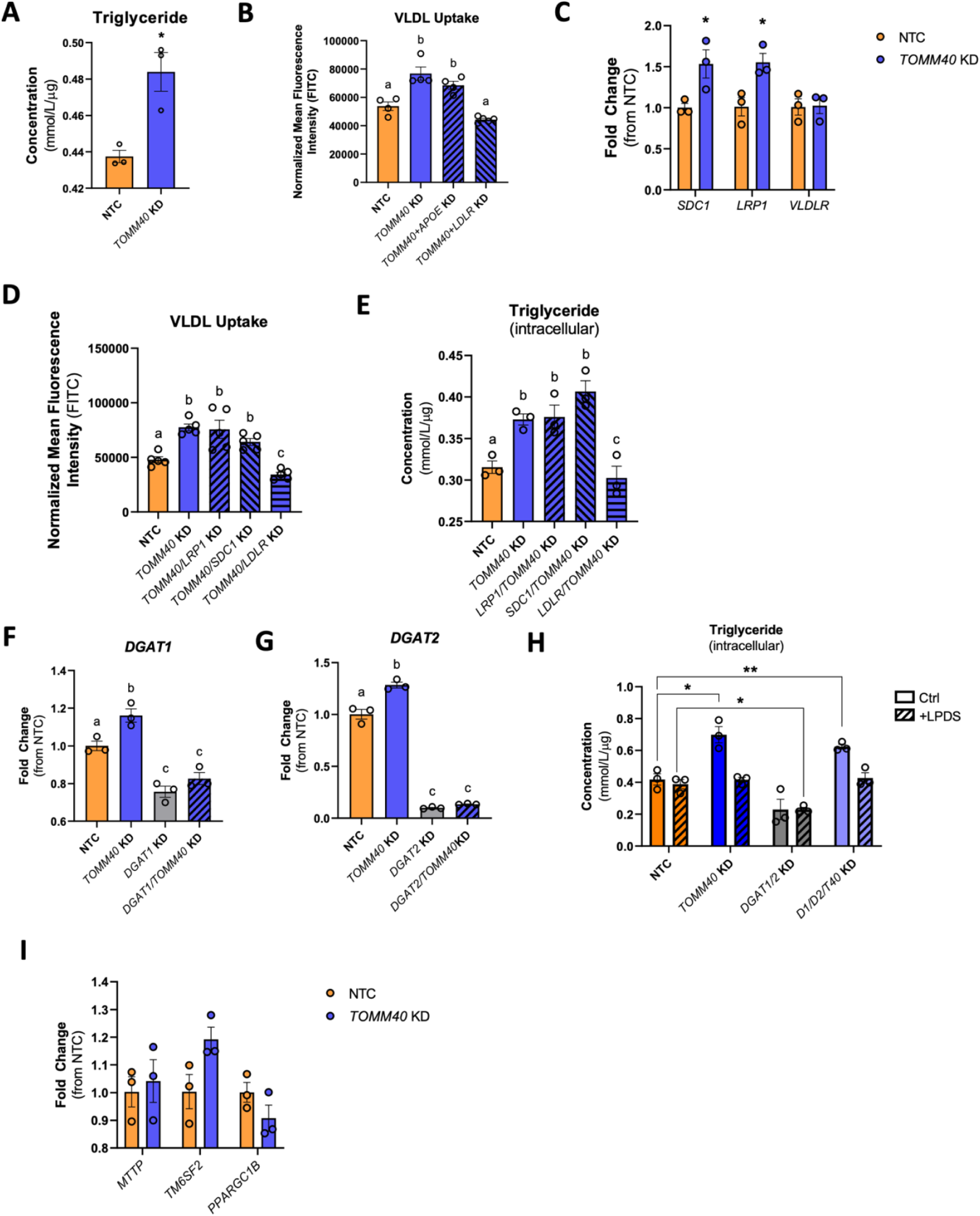
*TOMM40* KD promotes VLDL uptake and triglyceride accumulation via LDLR upregulation. (A) Quantification of intracellular triglyceride in HepG2 cells transfected with NTC vs. *TOMM40* siRNAs. (B) Quantification of DiI-labelled human VLDL uptake after 4 hr incubation by HepG2 cells transfected with siRNA for *TOMM40* singly and in combination with siRNAs for *APOE*, and *LDLR* KD, vs. NTC, measured by fluorescence spectrophotometry. (C) Relative *SDC1, LRP1*, and *VLDLR* mRNA levels in HepG2 cells transfected with NTC vs. *TOMM40* siRNAs, measured by qPCR. (D) Quantification of DiI-labelled VLDL uptake after 4 hr incubation by HepG2 cells transfected with siRNA for *TOMM40* singly and in combination with siRNAs for *LRP1, SDC1*, and *LDLR*, vs. NTC, measured by fluorescence spectrophotometry (E) Intracellular triglyceride levels quantified in HepG2 cells transfected with siRNA for *TOMM40* singly and in combination with siRNAs for *LRP1, SDC1*, and *LDLR,* vs. NTC. (F-G) Relative mRNA transcript levels of *DGAT1* and *DGAT2* in HepG2 cells transfected with NTC vs. siRNAs for *TOMM40, DGAT1*, and *DGAT2*, singly and in combination, measured by qPCR. (H) Quantification of intracellular triglyceride in *DGAT1/DGAT2* KD vs*. DGAT1/DGAT2/TOMM40* KD HepG2 cell incubated in 10% FBS or LPDS serum for 48 hrs. (I) Relative *MTTP, TM6SF2*, and *PPARGC1B* mRNA levels in HepG2 cells transfected with NTC vs. *TOMM40* siRNA, as measured by qPCR. For all: *p<0.05, **p<0.01, ***p<0.005, ****p<0.001 vs. NTC by one-way ANOVA, with post-hoc Student’s t-test to identify differences between groups. p<0.05 for *a* vs. *b* vs. *c* by two-way ANOVA, with Sidak’s multiple comparisons test. Data are represented as mean ± SEM. (*n=3* biological replicates)

We also considered the possibility that an increase in triglyceride synthesis may contribute to its cellular accumulation with *TOMM40* KD. However, KD of the genes encoding DGAT1 (**Fig 5F**) and DGAT2 (**Fig 5G**), enzymes with key roles in triglyceride synthesis, resulted in no significant reduction in triglyceride content of *TOMM40* KD cells (**Fig 5H**). Additionally, when cultured in lipoprotein deficient media, *TOMM40* KD HepG2 cells showed no increase in intracellular triglyceride content compared to NTC, confirming that VLDL uptake from the media plays a determining role in the accumulation of cellular triglyceride in HepG2 cells (**Fig 5H**). Finally, we showed that transcript levels of *MTTP, TM6SF2*, and *PPARGC1B,* which encode genes that have critical roles in regulating VLDL assembly and secretion^31,32^, were not increased by *TOMM40* KD (**Fig 5I**). We therefore conclude that increased VLDL uptake resulting from upregulation of *LDLR* is primarily responsible for the cellular triglyceride loading induced by *TOMM40* KD.

### Plasma cholesterol and triglyceride levels are reduced in AAV8-*Tomm40* shRNA C57BL/6J mice

We suppressed *Tomm40* expression with intraperitoneal (IP) injection of AAV8-*Tomm40* shRNA, which has strong tropism to liver cells^33^, in C57BL/6J male and female mice fed a western diet (0.15% cholesterol, 21% fat), achieving a KD efficiency in liver of greater than 60%. (**Fig 6A**). After 5 weeks, there were significant reductions in body weight that correlated with percent KD, however no changes were seen in food intake (**Fig S5**). TEM (transmission electron microscopy) of hepatic tissue sections from *Tomm40* KD male, but not female, mice demonstrated disruption of MERCs. (**Fig 6B-D, Fig S6**). Further results also consistent with those described above for *TOMM40* KD in HepG2 cells included reduced hepatic expression of *Mfn2* and increased expression of *Lxrb, Abca1, Cyp7a1, Srebf1c,* and *Ldlr* in both male and female mouse livers, though only males showed increased expression of *Apoe* (**Fig 6E,F**). Similar effects were observed in primary hepatocytes from 8-12-week old male mice transfected with *Tomm40* siRNAs *in vitro* (**Fig 6H**). Finally, plasma CA was higher in the *Tomm40* shRNA mice compared to controls though this was seen only in males (**Fig 6G**).

**Figure 6.**
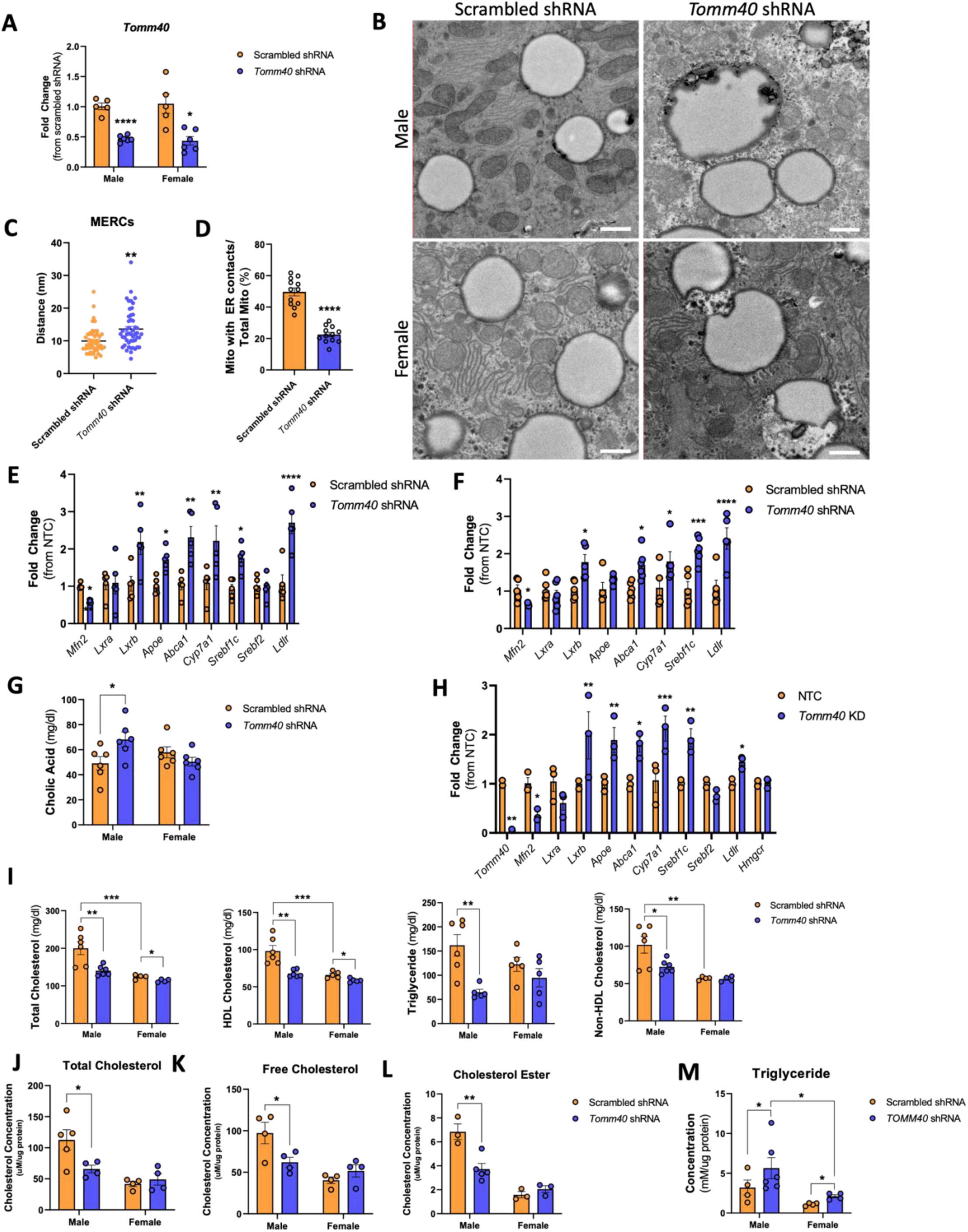
AAV8-T*omm40* shRNA injected C57BL/6J mice show reduced plasma cholesterol and triglyceride levels. (A) Relative mRNA transcript levels of *Tomm40* confirming KD in male and female mice of ∼60%, quantified by qPCR. (n=5-6 mice/sex/group) (B) Representative TEM images of scrambled shRNA (control) vs. *Tomm40* shRNA in male and female, mice. Scale bars, 1μm. (C-D) Analysis of MERCs using ImageJ software: (C) ER-mitochondria distance (nm), (N) length of MERCs (nm), (D) percentage of mitochondria with ER contacts out of total mitochondria per cell. (*n= 12-48* fields) (E-F) Relative mRNA transcript levels comparing scrambled vs. *Tomm40* shRNA mice in male (E) and female (F). (*n=4-6* mice/sex/group) (G) Quantification of plasma cholic acid levels in scrambled vs. *Tomm40* shRNA, male and female mice, measured by ELISA. (*n=4-6* mice/sex/group) (H) Relative mRNA transcript levels comparing NTC vs. *Tomm40* siRNA transfected primary hepatocytes derived from male C57BL/6J mice, quantified by qPCR. (*n=3* male mice/group) (I) Plasma total cholesterol, HDL-cholesterol, and triglyceride from male and female mice were quantified with AMS Liasys 330 Clinical Chemistry Analyzer, and Non-HDL-cholesterol was calculated by subtracting HDL-cholesterol from total cholesterol levels. (*n=5-6* mice/sex/group) (J-L) Hepatic total cholesterol (J), free cholesterol (K), and cholesterol ester (L) levels quantified from male and female mice liver, using Amplex Red Cholesterol Assay. (*n=4-6* mice/sex/group) (M) Quantification of hepatic triglyceride levels in male and female mice livers, using EnzyChrom™ Triglyceride Assay. (*n=4-6* mice/sex/group) For all: *p<0.05, **p<0.01, ***p<0.005, ****p<0.001 vs. NTC by one-way ANOVA, with post-hoc Student’s t-test to identify differences between groups. Data are represented as mean ± SEM.

*In vivo Tomm40* KD resulted in substantially lower plasma concentrations of total cholesterol, non-HDL cholesterol, triglycerides, and HDL cholesterol, albeit to a lesser extent in female than in male mice, in which levels of all these measures with scrambled shRNA treatment were significantly higher than in females. (**Fig 6I**). These findings were supported by measurements of lipoprotein particle concentrations using ion mobility methodology (**Fig S7**). Notably, we also found decreased hepatic cholesterol content (**Fig 6J-L**) in male mice, and increased triglyceride content (**Fig 6M**) in both sexes, as was seen in our *in vitro* experiments in HepG2 cells.

### *TOMM*40 KD induces lipid droplet accumulation and metabolic-dysfunction associated steatotic liver disease (MASLD)

Consistent with the increased triglyceride content of *TOMM40* KD HepG2 cells, TEM micrographs revealed an increase in lipid droplet (LD) number and size (**Fig 7B,C**), and further cytometric examination showed greater Nile red fluorescence than in NTC treated cells (**Fig 7D,E**). With double knockdown of *TOMM40* and either *APOE* or *LDLR* in HepG2 cells, there was a significant reduction in Nile red fluorescence vs. *TOMM40* KD alone (**Fig 7E**), suggesting that the accumulation of LDs with *TOMM40* KD is in part due to lipoprotein uptake via upregulation of the APOE/LDLR pathway. Similar to the findings in HepG2 cells, Oil Red O (ORO) staining of livers from both male and female mice following *in vivo Tomm40* KD demonstrated a greater number of LDs vs. the scrambled shRNA controls (**Fig 7H-J, S8**). Consistent with these findings, significant hepatic steatosis in both sexes was confirmed by hematoxylin-eosin staining of liver sections (**Fig 7S-T**), as well as increased liver-to-bodyweight ratio (**Fig 7U**), and elevated plasma AST (**Fig 7V**), though plasma ALT was increased only in males (**Fig 7W**).

**Figure 7.**
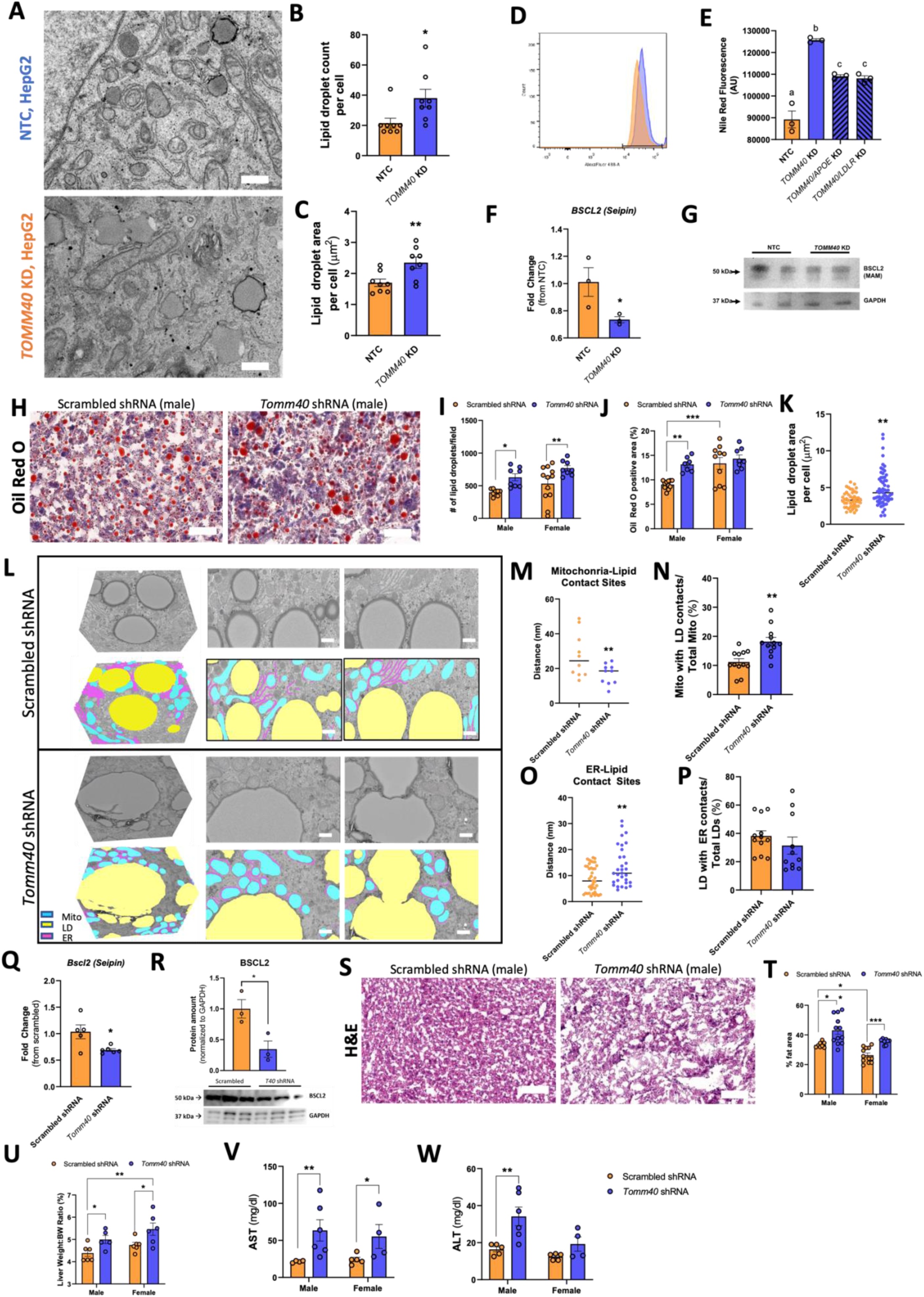
*TOMM40/Tomm40* KD induces lipid droplet accumulation and hepatic steatosis *in vivo*. (A)Representative TEM images of lipid droplets in NTC vs. *TOMM40* KD HepG2 cells. Scale bars, 1000 nm. (B-C) Analysis of lipid droplets in HepG2 cells using ImageJ software: (B) lipid droplet count per cell, (C) average lipid droplet area per cell (µm^2^). (*n= 8-12* cells) (D) Representative flow cytometry histogram of NTC vs. *TOMM40* KD HepG2 cells. Neutral lipids were stained with Nile red. (E) Analysis of flow cytometry data in NTC vs. *TOMM40* KD, singly and in combination with *APOE* and *LDLR* KD, HepG2 cells stained with Nile red. (*n=3* biological replicates) (F) Relative mRNA transcript levels of *BSCL2* (Seipin) in NTC vs. *TOMM40* KD HepG2 cells. (*n=3* biological replicates) (G) Representative western blot of BSCL2 protein expression in isolated MAMs of NTC vs. *TOMM40* KD HepG2 cells. (H) Representative ORO images of scrambled vs. *Tomm40* shRNA male mouse liver (magnification x 400). Scale bars, 50 µm. (I-J) Analysis of ORO staining using ImageJ software: (I) # of lipid droplets/field, (J) ORO positive area (%). (*n=10-15* fields/sex/group) (K) Analysis of lipid droplet surface area (µm^2^) in male mouse liver tissue using ImageJ software. (*n=24-48* fields) (L) Representative FIB-SEM micrographs of 3D reconstructed and 2D slices from scrambled vs. *Tomm40* shRNA male mice liver. Segmentation analysis indicates mitochondria (blue), lipid droplets (yellow), and endoplasmic reticulum (purple). Scale bars, 1 µm. (M-P) Analysis of organelle contact sites in scrambled vs. *Tomm40* shRNA male mice liver from TEM images using ImageJ software: (M) mitochondria-lipid contact site distance (nm), (N) percentage of mitochondria with lipid droplet contacts out of total mitochondria (%), (O) ER-lipid contact site distance (nm), (P) percentage of lipid droplets with an ER contact site out of total lipid droplets (%). (*n=10-40* fields) (Q) Relative mRNA transcript levels of *Bscl2* (Seipin) in scrambled vs. *Tomm40* shRNA male mice livers. (*n=5-6* mice/group) (R) Relative protein amount and representative western blot of BSCL2 protein expression in isolated MAMs of scrambled vs. *Tomm40* shRNA male mice livers. (S) Representative hematoxylin-eosin stained images of scrambled vs. *Tomm40* shRNA male mouse livers (magnification x 400). Scale bars, 50 µm. (T) Analysis of % fat area/field in hematoxylin-eosin stained liver samples using ImageJ software. (*n=10-15* fields/sex/group) (U) Liver weight-to-body weight ratio (%) of scrambled vs. *Tomm40* shRNA mice. (*n=5-6* mice/sex/group) (V-W) Plasma AST and ALT from male and female mice were quantified with AMS Liasys 330 Clinical Chemistry Analyzer. (*n=5-6* mice/sex/group) For all: *p<0.05, **p<0.01, ***p<0.005, ****p<0.001 vs. NTC by one-way ANOVA, with post-hoc Student’s t-test to identify differences between groups. Data are represented as mean ± SEM.

Although *Tomm40* KD in both sexes induced hepatic steatosis, the intracellular mechanisms were found to be different. Imaging the 3D morphology of LDs at MERCs by focused ion beam scanning electron microscopy (FIB-EM) (**Fig 7L**) in conjunction with TEM analysis revealed not only an increase in average LD surface area per cell (**Fig 7K**), but also increased LD-mitochondria contact sites (**Fig 7M-N**) and reduced LD-ER contact sites in the livers of male (**Fig 7O-P**), though not in female (**Fig S9),** *Tomm40* KD mice. The evidence for increased LD formation from mitochondria rather than the ER in males was supported by reduced expression of *Bscl2*, the gene encoding seipin, which is known to play a key role in LD formation from the ER at MERCs^34^ (**Fig 7Q**). Moreover, in isolated hepatic mitochondria-associated membranes (MAMs) containing MERCs, we consistently observed reduced BSCL2 protein expression in males (**Fig 7R**). Having confirmed the reduction in *BSCL2* gene and protein expression (MAMs fraction) in *TOMM40* KD HepG2 cells (**Fig 7F,G**), we showed this effect to be LXR-independent by treatment with GSK2033 (**Fig S11**). In female mice livers, *Tomm40* KD did not affect BSCL2 protein expression in MAM fractions (**Fig S10**), which may explain the lack of disruption of LD-ER contact sites. No changes in cytosolic BSCL2 protein levels were found in either sex (**Fig S10**). Despite sex-differences, these results suggest that decreased BSCL2-mediated LD formation from the ER with *TOMM40* KD to play a potential auxiliary role, specifically in males, in the uptake of triglyceride-rich lipoproteins via LDLR.

## DISCUSSION

TOMM40 is a subunit of the TOM complex that has a key role in mediating uptake of mitochondria-targeted proteins, maintaining mitochondrial membrane potential, and reducing the mitochondrial stress response^14,15^. We show here, by KD of *TOMM40* in hepatocytes and *in vivo* in mice, that TOMM40 also has multiple effects that impact hepatic lipid and plasma lipoprotein metabolism. A primary mechanism for these effects is increased activation of LXRs as well as greater *LXRB* gene expression with *TOMM40* KD that resulted in increased transcription of multiple genes affecting cholesterol and lipoprotein metabolism. Notably, we found that LXR-mediated upregulation of both *APOE* and *LDLR* with *TOMM40* KD led to increased hepatocellular uptake of LDL as well as VLDL particles. We further demonstrated by concurrent KD of *MFN2* that disruption of MERCs was a major mediator of this LXRB-specific process, in conjunction with increased ROS and generation of oxysterols, in particular ROS-derived 7-ketocholesterol, a known ligand of LXR^35,36^. Interestingly however, *TOMM40* KD but not *MFN2* KD increased levels of oxysterols that are produced by enzymatic rather than oxidative reactions, suggesting that effects other than MERC disruption and mitochondrial dysfunction may also contribute to LXR activation by *TOMM40* KD. Other than 7-ketocholesterol, the increase in *CYP3A4* expression, the rate limiting step for 4β-OHC, in both *MFN2* KD and *TOMM40* KD HepG2 cells, could be explained by activation of pregnane X receptor (PXR), another nuclear receptor and transcriptional regulator of CYP3A4, that may be due to MERC-induced cellular stress response^37,38^. Furthermore, increased *CYP3A4* expression and thus 4β-OHC have been shown to regulate lipogenic genes, including *SREBP1c,* and liver triglyceride levels via LXR activation^39^. However, future experimentation is required to confirm the role of 4β-OHC in mediating effects of *TOMM40* KD.

Notably, despite demonstrating that *TOMM40* KD increased cellular uptake of LDL via upregulation of LDLR and APOE, intracellular cholesterol levels were reduced, a finding confirmed in male mouse livers *in vivo*. This was shown to be primarily due to increased expression of *ABCA1*, a LXR transcriptional target^40^, in that KD of *ABCA1* abrogated this effect of *TOMM40* KD. Reduction in cellular cholesterol ester content could also have been impacted by the observed reduced expression of *ACAT1*, the enzyme responsible for the esterification of free cholesterol^41^ that is enriched at MERCs^42^.

It has been reported that LXR, specifically LXRA, can directly regulate *LDLR* gene expression in hepatoblastoma cells^43^. We have further shown that LXR can induce increased expression of *SREBF1c*^44^, and our finding that KD of this gene abrogated the increase in *LDLR* expression with *TOMM40* KD points to its upregulation by LXR in mediating the *LDLR* response, likely by binding to the *LDLR* promoter region as shown from our luciferase experiment and previously in HepG2 cells^45,46^. While SREBF2 is the major transcription factor regulating increased expression of LDLR in response to reduced cellular unesterified cholesterol content^47^, our finding that expression of *HMGCR*, a canonical SREBF2 target^48^, is not affected by *TOMM40* KD suggests that SREBF2 activity does not play a significant role in *TOMM40* KD-induced LDLR upregulation.

Among the genes upregulated by *TOMM40* KD in HepG2 cells were *CYP7A1* and *CYP27A1*, which mediate the rate-limiting steps in the synthesis of the bile acids cholic acid (CA) and chenodeoxycholic acid (CDCA), respectively^49,50^. Consistent with the *CYP7A1* effect, we found that both intracellular levels and secretion of CA were increased by *TOMM40* KD, representing another route for disposing of cellular cholesterol. While *Tomm40* KD male mice showed increased plasma CA levels, no clear differences were detected in females. These results can be explained by previously identified sex-linked differences in not only hepatic bile acid production and secretion^51^, but also composition, storage, and release of bile acids from the gallbladder^52^ and bile acid metabolism in the gut microbiota^53^, all known to contribute to plasma CA levels independent of CYP7A1 activity. Furthermore, it has been reported that women have lower circulating bile acid concentrations than men, which may protect from development of hepatocellular carcinoma^54^. In contrast, despite the increased expression of *CYP27A1*, independent of LXR activity (**Fig S12**), cellular CDCA content was reduced. Synthesis of the CDCA precursor 27-OHC occurs in mitochondria^55,56^, separate from LXR regulation, and the interaction of TOMM40 with STAR at MERCs has been shown to promote transport of cholesterol into mitochondria^8,57^. We observed that *STAR* expression is reduced by *TOMM40* KD, and suggest that this, together with MERC disruption as shown by *MFN2* KD, leads to a decrease of mitochondrial cholesterol that is available for 27-OHC and CDCA synthesis.

In contrast to the reduction of cholesterol in HepG2 cells by *TOMM40* KD, we observed a significant increase in cellular content of triglycerides. This is likely attributable to stimulation of uptake of triglyceride-rich VLDL, which as for LDL, was shown to be mediated by upregulation of *LDLR* in conjunction with increased expression of *APOE*. While *TOMM40* KD also increased expression of genes encoding other APOE-binding proteins - *LRP* and *SDC1*^58^ - KD of these genes had no effect on VLDL uptake. Moreover, there was no phenotypic effect of hepatic *TOMM40* KD on expression of genes regulating VLDL secretion or triglyceride synthesis. The increase in cellular triglyceride was manifest by greater number and size of LDs, and the dependence of their formation on *LDLR*-mediated triglyceride uptake was confirmed by demonstrating reduced lipid staining with KD of either *LDLR* or *APOE*. It has previously been shown in astrocytes that APOE can also act as a LD surface protein regulating LD size and triglyceride saturation, independent of an effect at the ER lumen^59^.

Consistent with the cellular findings, *in vivo* KD of *Tomm40* resulted in increased hepatic triglyceride and reduced cholesterol content, and increased size and number of hepatic LDs. Consequently, *Tomm40* KD mice developed hepatic steatosis based on histological (H&E and ORO) assessments and quantification of plasma AST and ALT levels. Apart from LD accumulation in hepatocytes, the remodeling of LD contact sites and dynamics in response to metabolic changes in the liver is another defining characteristic of hepatic steatosis and MASLD^60^. In this regard, we have shown that *TOMM40* KD reduced LD-ER contact sites in HepG2 cells via a reduction in BSCL2 (seipin) expression in MAMs containing MERCs. Loss of BSCL2 at LD-ER interfaces has been shown to result in abnormal LDs including supersized variants, consistent with our findings^61,62^. By TEM we have shown that in conjunction with reduction of LD-ER contact sites there was an increase in LD-associated mitochondria with *TOMM40* KD, an effect that may lead to increased trafficking of triglyceride-derived fatty acids via LDs into the mitochondria^63^ as well as the upregulation of fatty acid oxidation that has been associated with MASLD^64^.

While the effects of *TOMM40 KD* on LD-ER and LD-mitochondria contact sites and BSCL2 (seipin) expression in HepG2 cells were replicated in liver tissue from male mice following in vivo *Tomm40* KD, these effects were not observed in females. Thus, mechanisms other than suppression of the seipin pathway are responsible for promoting LD accumulation and hepatic steatosis with *Tomm40* KD in females, and it is possible that these mechanisms are also operative in males. For example, mediators of LD formation located outside of MERCs, including FITM2, regulate the budding of LDs from ER into the cytosol^65^, and coalescence of LDs is promoted by the CIDE family of proteins (aka FSP27)^66,67^. Additionally, it is notable that sex hormones including estrogen, have been reported to transcriptionally regulate genes involved in mitochondrial dynamics and mitophagy, including *MFN2* expression, which may further explain the differential effects of *Tomm40* KD on MERCs and mito-LD contact sites between male and female mouse livers^68,69^.

In addition to promoting hepatic steatosis, *in vivo* KD of *Tomm40* in mice resulted in significant reductions in plasma lipid and lipoprotein levels that were greater in males. This may reflect sex differences in lipoprotein production and/or clearance^70^ that resulted in higher baseline plasma lipid levels in the males^71^, as well as differing age-related effects^72,73^ and the use of a western diet, since, for example, female C57BL/6J mice are resistant to high fat diet-induced obesity^74^.

While a strength of this study is the identification of significant effects of hepatic *TOMM40/Tomm40* KD on lipid and lipoprotein metabolism in both a human liver cell and an *in vivo* mouse model, further study will be required to replicate the findings in other hepatic cell lines and mouse strains, and to test the *in vivo* effects in conjunction with other dietary interventions, especially in light of the sex difference in lipid metabolism noted above and the report that high-fat diets promote formation of LD contact sites with mitochondria and ER^63^. Finally, kinetic studies of the effects *Tomm40* KD on lipoprotein production and clearance are needed to determine the basis for the reductions in their plasma levels and the differing effects in male and female mice. In this regard, it remains possible that extrahepatic *Tomm40* KD with our AAV8-shRNA, as we have observed in skeletal muscle^75^, could have contributed to these plasma lipid changes.

Our findings suggest that identification of agents which increase hepatic TOMM40 expression, and in doing so, maintain mitochondrial function and MERCs, could provide new therapeutic opportunities for MASLD, as well as other conditions such as Alzheimer’s disease in which mitochondrial pathology plays a role. Finally, our discovery that *TOMM40* KD increases expression of the adjacent *APOE* gene suggests a novel connection between mitochondrial function and lipid metabolism.

## METHODS

### Mice studies

6-week old C57BL/6J male and female mice (*n=6* per group) were purchased from Jackson Laboratory (Bar Harbor, ME) and placed on a western diet (0.15% cholesterol, 21% fat, D12079Bi, Research Diets). At 8 weeks of age, mice were intraperitoneal (IP) injected with either 4×10^11^ GC AAV8-Tomm40 shRNA or AAV8-CMV-null as a control (VectorBuilder). Weekly bodyweight and food intake measurements were recorded. At 14-weeks old, unfasted mice were terminated and liver tissues were collected, snap frozen in liquid nitrogen, and immediately transferred to −80°C freezer. For electron microscopy, tissues were immediately fixed in 2% glutaraldehyde + 2% paraformaldehyde solution after termination. Blood was collected by cardiac puncture and plasma separated via centrifugation at 850 x g for 15 min at 4°C. All experiments were done blinded and randomized. Animal research was approved by the University of California, San Francisco, Laboratory Animal Resource Center.

### Primary mouse hepatocytes

Hepatocytes were isolated from wild-type, 8-12 week-old C5BL/6J male mice (Jackson Laboratory) by the UCSF Liver Center according to the protocol established by Desai et al.^80^ Hepatocytes were cultured in DMEM with 5% fetal calf serum (Hyclone), L-glutamine, penicillin-streptomycin antibiotic, insulin-transferrin-selenium, and HEPES (Gibco). *Tomm40* and non-targeting control (NTC) siRNAs were transfected into the primary hepatocytes using TransIT-TKO® transfection reagent (Mirus Bio) according to the manufacturer’s protocol for 48 hr after hepatocyte plating at 37°C and 5% CO_2_.

### HepG2 cell culture

HepG2 human hepatoma cells were grown in EMEM (Eagle’s Minimum Essential Medium; ATCC) supplemented with 10% fetal bovine serum (FBS; Thermo Fisher Scientific) or 10% lipoprotein-deficient serum (LPDS; Thermo Fisher Scientific), and 1% penicillin-streptomycin (Gibco) at 37°C and 5% CO_2_. Cells were passaged every 7 days. Cells were routinely tested for mycoplasma using MycoAlert™ PLUS mycoplasma detection kit (Lonza) and only mycoplasma negative cells were used.

Knock-down (KD) of *TOMM40, MFN2, LXRA, LXRB, APOE, LDLR, DGAT1, DGAT2, LRP1, SDC1, ABCA1*, and *SREBF1*, was achieved by addition of their respective siRNAs (10 μM) using Lipofectamine RNAiMAX transfection reagent (Life Technologies) and Opti-MEM I (Gibco) according to the manufacturer’s instructions for 48 hrs.

Human pCMV-expressing-TOMM40 (NM_001128916.2), STAR (NM_000349.3), and empty vector (EV; ORF_stuffer) plasmids stored in bacterial glycerol stocks were purchased from VectorBuilder Inc. Expression plasmids were cultured on Luria-Bertani (LB) Agar plates containing ampicillin at 37°C. Single colonies were selected and grown separately in LB broth at 37°C with continuous shaking (225 rpm) overnight. DNA plasmids were extracted by ZymoPURE II Plasmid Midiprep Kit (Zymogen) according to the manufacturer’s protocol. Cells were transiently transfected with purified plasmids using Lipofectamine 3000 transfection reagent (Thermo Fisher Scientific).

### Mitochondrial respiration measurements

HepG2 cells were seeded at 2,000 per well in 96-well plates with XF assay medium (Agilent) supplemented with 2 mM sodium pyruvate (Gibco), 2 mM GlutaMAX™ (Gibco), and 10 mM glucose (Sigma), at pH 7.4. During experimentation, 1.5 µM oligomycin, 2 µM FCCP, and 2 µM Antimycin A + Rotenone (Seahorse XF Cell Mito Stress Test Kit, Agilent) were added sequentially to the Agilent Seahorse XFe96 Extracellular Flux Analyzer via injection ports to determine basal and maximum respiration, ATP production, and proton leak. Oxygen consumption rate (OCR) values were presented with non-mitochondrial oxygen consumption deducted and normalized to total protein concentration per well using BCA assay (Genesee Scientific).

### Mitochondrial assays

Cellular reactive oxygen species (ROS) were quantified by the DFCDA/H2DCFDA – Cellular ROS Assay Kit (Abcam) according to the manufacturer’s instructions. Cells were stained with DFCDA for 45 min and washed in 1X DPBS. Fluorescence intensity was measured by a fluorescence spectrophotometer with excitation/emission at 485 nm/535 nm.

### Calcium imaging

Mag-Fluo-4AM was used to quantify free Ca2+ within the ER lumen^81^. HepG2 cells were seeded in a black clear-bottom 96-well plate and transfected with siRNAs for 48 hrs. Cells were then washed with Hank’s Buffered Saline Solution (HBSS) containing 20 mM Hepes, and then incubated with 10 µM Mag-Fluo-4 AM (Thermo Fisher Scientific) for 1 hr at 37°C in HEPES-buffered saline (135 mM NaCl, 5.9 mM KCl, 11.6 mM HEPES, 1.5 mM CaCl2, 11.5 mM glucose, 1.2 mM MgCl2, pH 7.3), supplemented with BSA (1 mg/mL) and pluronic acid (2%, v/v). Excess dye was then washed off using HBSS and measured on a fluorescent spectrophotometer at a wavelength of 495/515 (Ex/Em).

### TEM sample preparation

HepG2 cells were grown on MatTek glass bottom dishes (P35G-1.5-14-C, MatTek) and fixed in 2% glutaraldehyde + 2% paraformaldehyde solution (prepared by Electron Microscopy Lab, UC Berkeley) for 24 hrs. Cells and tissues were post-fixed in 1% osmium tetroxide in 0.1 M sodium cacodylate buffer, pH 7.2, for 1-2 hrs. Cells were dehydrated in a serial diluted ethanol solution from 30 to 100%. Liver tissues were embedded in increasing concentrations of Durcupan resin. HepG2 cells for TEM imaging were infiltrated with 50% Epon-Araldite resin (containing benzyldimethylamine (BDMA) accelerator), followed by 100% resin for 1 hr each. Cells and tissues were polymerized at 60C for 24hrs.

### Transmission electron microscopy

The resin embedded sample blocks were trimmed, and 90 nm ultrathin sections were cut using a Leica UC6 ultramicrotome (Leica Microsystems, Vienna, Austria) and collected onto formvar-coated slot grids. Sections were imaged to find target regions using a Tecnai 12 120kV TEM (FEI, Hillsboro, OR, USA) and data recorded using an Gatan Rio16 CMOS camera and GMS3 software (Gatan Inc., Pleasanton, CA, USA).

### Volume imaging processing for liver tissue

200 µm thick slices from previously fixed material were stained using an osmium-thiocarbohydrazide-osmium (OTO) method^82,83^ in combination with microwave-assisted processing, followed by high pressure freezing and freeze substitution (HPF-FS), as previous described by Ewald et al.^84^ Briefly, samples were OTO stained, incubated with 2% aqueous uranyl acetate overnight, subjected to HPF followed by super quick FS^85^ with 4% osmium tetroxide, 0.1% uranyl acetate and 5% ddH2O in acetone, and embedded and polymerized in hard epon resin.

### Focused ion beam scanning electron microscopy (FIB-SEM) imaging

The trimmed sample blocks were glued with silver paint (Ted Pella Inc.) onto Al stubs, and sputter coated (Pd/Au) with a Tousimis sputter coater on top of a Bio-Rad E5400 controller. Focused Ion Beam Scanning Electron Microscopy (FIB-SEM) imaging was performed using a Zeiss Crossbeam 550 (Carl Zeiss Microsystems GmbH, Oberkochen, Germany). The sample was tilted at 54° in order to be perpendicular to the ion beam. FIB milling and SEM imaging of the target area were set up using Atlas 5 3D tomography (Carl Zeiss Microsystems GmbH, Oberkochen, Germany). Slices with a thickness of 10 nm were milled from the target area using a 30 kV 300 pA ion beam. Energy-selective Backscattered (ESB) images were collected at 1.5 kV 1nA with a dwell time of 18 ns, image pixel size of 10 nm, and tilt correction angle of 36°.

### Image processing

TEM and FIB-SEM images were analyzed using ImageJ software according to the method of Lam et al. Contact sites between organelles were identified manually from high-resolution, high-magnification TEM images, and outlined using an optical pen in ImageJ to calculate structural parameters. MERCs (and other contact sites) were identified by having a gap of 10-30 nm between the outer mitochondrial membrane, the ER, or the lipid droplet membrane, as well as having the ER be ribosome-free^86^.

The collected FIB-SEM images were aligned with the Slice Registration in Dragonfly 2022.2 (Comet Technologies Canada Inc., Canada). The Dragonfly Segmentation Wizard and Deep Learning Tool were used to segment organelles (lipid droplets, mitochondria, endoplasmic reticulum (ER)).

### Enzyme-linked immunosorbent assay (ELISA)

Cells, supernatant, and liver tissues were lysed in M Cellytic Lysis Buffer containing 1% protease inhibitor (Halt™ Protease Inhibitor Cocktail; ThermoFisher Scientific) for 15 min using a cell disruptor or homogenizer. The lysate was centrifuged at 14,000 x g for 15 min and supernatant was collected. Oxysterols, 25-, 24(S)-, and 27-OHC, were quantified by ELISA kits purchased from MyBioSource according to the manufacturer’s protocol. Total bile acid, cholic acid, and chenodeoxycholic acid levels were measured in cells and cell media (supernatant), by ELISAs purchased from Cell Biolabs Inc, according to the manufacturer’s protocol. In addition, cholic acid was quantified in mouse plasma samples using the same kit from Cell Biolabs Inc. Samples from each experiment were normalized to protein levels quantified by BSA assay (Genesee Scientific).

### Fluorescence imaging

To quantify LDL and VLDL uptake, HepG2 cells were loaded with 10 µM fluorescently labelled BODIPY™ FL LDL (Thermo Fisher Scientific) or DiI-labelled VLDL (Kalen Biomedical LLC) isolated from human plasma, for 4 hrs at 37°C. Cells were washed and reconstituted in 1X PBS buffer and read on a fluorescence spectrophotometer. Fluorescence readings were normalized to total protein concentration per well by BCA assay (Genesee Scientific).

LDLR cell surface protein was quantified by fixing cells in 4% paraformaldehyde and incubated with anti-LDLR at 1:100 (Santa Cruz Biotechnology, sc18823) for 45 min, followed by goat anti-mouse IgG-FITC at 1:400 (Santa Cruz Biotechnology, sc-2010) for 30 min. Intracellular 7-ketocholesterol was measured by fixing cells in 4% paraformaldehyde and staining with 7-ketocholesterol monoclonal antibody at 1:50 (3F7, Invitrogen, MA5-27561) diluted in 1X permeabilization buffer (Invitrogen) for 1 hr, followed by goat anti-mouse IgG-FITC at 1:400 (Santa Cruz Biotechnology, sc-2010) for 30 min.

To quantify lipid droplets, cells were stained with 100 µM Nile Red (Sigma) for 30 min. Fluorescence for all experiments described in this section was quantified on a BD LSRFortessa™ Cell Analyzer flow cytometer with a median fluorescence value of 10,000 gated events. Data was analyzed using FlowJo v10.7.1.

### Lipid extraction and quantification

Mouse liver samples were homogenized by GentleMACS™ dissociator (Miltenyi Biotec) and cells were dissociated using a cell disruptor (Bio-Rad) in a mixture of chloroform, methanol, and water (8:4:3, v/v/v) (ref) or hexane-isopropanol (3:2, v/v) for lipid and cholesterol extraction, respectively. Hepatic and intracellular triglyceride (TAG) and glycerol were quantified with EnzyChrom™ Triglyceride or Glycerol Assay Kit (BioAssay Systems) according to the manufacturer’s protocol. Lipid concentrations were measured at an absorbance of 570 nm. Cholesterol samples were dried under nitrogen gas and reconstituted with buffer (0.5 M potassium phosphate, pH 7.4, 0.25 M NaCl, 25 mM cholic acid, 0.5% Triton X-100). Intracellular cholesterol levels were then quantified with the Amplex Red Cholesterol Assay Kit (Life Technologies) according to the manufacturer’s protocol.

### Isolation of mitochondria and mitochondria-associated membranes

Mitochondria-associated membranes (MAMs) were isolated from HepG2 cells and liver according to the method of Wieckowski et al.^87^ Cells were rinsed in 1X PBS twice, dislodged with 0.25% Trypsin-EDTA (Gibco), washed again in 1X PBS and centrifuged at 600 x g for 5 min at 4°C. Pelleted cells and liver tissue were resuspended in MSHE + BSA buffer (210 mM mannitol, 70 mM sucrose, 5 mM HEPES, 1 mM EGTA, and 0.5% BSA, at 7.2 pH). Samples were transferred to a small glass dounce and homogenized. The homogenate was centrifuged at 600 x g for 10 min at 4°C, and the supernatant was extracted and centrifuged at 8,000 x g for 10 min at 4°C. The pellet containing the isolated crude mitochondria was used to quantify mitochondrial cholesterol content. Furthermore, the isolated crude mitochondria were resuspended in MSHE + BSA buffer and percoll medium (Gibco) and centrifuged in a Beckman Coulter Optima L-100 XP Ultracentrifuge (SW40 rotor, Beckman) for 95,000 x g for 30 min at 4°C. The white band located above the mitochondria pellet was identified as the MAM fraction and collected and diluted ten times with MSHE + BSA buffer. The fraction was centrifuged at 100,000 x g for 1 hr (70-Ti rotor, Beckman) at 4°C to isolate the MAMs for immunoblotting.

### Immunoblotting

Cells and liver tissues were lysed in M Cellytic Lysis Buffer containing 1% protease inhibitor (Halt™ Protease Inhibitor Cocktail; ThermoFisher Scientific) for 15 min using a cell disruptor or homogenizer, respectively. The lysate was centrifuged at 14,000 x g for 15 min, supernatant was collected, and protein concentration was measured by BCA assay (Genessee Scientific). Proteins were separated on a 4-20% Tris-polyacrylamide gradient gel (Bio-Rad) and transferred onto a nitrocellulose membrane using the iBlot™ 2 Gel Transfer Device (ThermoFisher Scientific). Membranes were blocked in Tris-buffered saline with 0.1% tween (TBST) + 5% milk for 2 hrs to minimize non-specific antibody binding. Membranes were then incubated with primary antibodies diluted 1:1000 (v/v) in TBST overnight on a rotating platform at 4°C. After washing in TBST, membranes were incubated with secondary antibodies, anti-rabbit IgG (7074) and anti-mouse IgG (7076), HRP-linked antibodies (Cell Signal) at 1:2500 (v/v) dilution, for 30 min before a last series of washes. SuperSignal™ West Pico PLUS Chemiluminescent Substrate (ThermoFisher Scientific) was added to the membrane to visualize proteins (30). All antibodies used are listed in the key resources table.

### RT-qPCR

RNA was extracted from liver tissue and cell samples using RNeasy Mini Qiacube Kit (Qiagen) with the Qiacube Connect (Qiagen) according to the manufacturer’s protocol. cDNA synthesis from total RNA was performed using the High Capacity cDNA Reverse Transcription Kit (Applied Biosystems). Primers were designed and obtained from Elim Biopharmaceuticals and run with SYBR™ Green qPCR Master Mix (Thermo Fisher Scientific) on an ABI PRISM 7900 Sequence Detection System to quantify mRNA transcript levels. RT-qPCR primers used in this study are listed in the key resources table. The mean value of triplicates for each sample was normalized to GAPDH (human) or 18s (mouse) as the housekeeping genes.

To quantify mRNA stability, Actinomycin D (Life Technologies) at a final concentration of 1 µg/mL was added to HepG2 cells treated with siRNAs at varying time intervals between 0-240 mins. At each time point, qPCR was conducted to quantify *LDLR* mRNA transcript levels. The half-life of the *LDLR* mRNA transcript in HepG2 cells with *TOMM40* KD was compared to the NTC group.

### Dual luciferase reporter assay

The dual luciferase assays were performed as previously described in Smith et al.^88^ HepG2 cells were seeded on 96-well plates with 100 µL of low glucose EMEM + 10% FBS or 10% LPDS (lipoprotein-deficient serum). On day of seeding, cells were transfected with indicated siRNAs. After 48 hrs, cells were transfected with indicated 100 ng LDLR luciferase construct + 1 ng secreted nanoluciferase construct in a total of 10 μL OptiMEM using Lipofectamine 3000 transfection reagent (Life Technologies) according to the manufacturer’s instructions. 48 hours post LDLR-luciferase transfection and treatment, 10 µL of media from each well was added to 10 µL of non-lytic 2x coelenterazine reagent (300 mM sodium ascorbate, 5 mM NaCl, 0.1% BSA, 40 μM coelenterazine (Goldbio CZ25)) in separate wells of a 384-well plate. Nanoluciferase plate was shielded from light incubated on a shaker at room temperature for 10 mins. Firefly luciferase activity was immediately evaluated in the plates containing the cells by adding 100 µL of 2x firefly lytic assay buffer (100 mM Tris-HCl pH 7.7, 50 mM NaCl, 2 mM MgCl2, 0.17% Triton X-100, 10 mM DTT, 0.4 mM coenzyme A, 0.3 mM ATP, and 0.28 mg/ml luciferin (Goldbio LUCK-1G)). Raw luminescence for all plates was measured on a plate reader with 1 second integration time. Readout of firefly luciferase in each well was normalized to the corresponding secreted nanoluciferase control.

### Plasma lipid and lipoprotein analyses

Total cholesterol (TC), HDL-C, triglyceride, AST (aspartate aminotransferase) and ALT (alanine transaminase) levels were measure by enzymatic end-point measurements using enzyme reagent kits (Kamiya Biomedical) in an AMS Liasys 330 Clinical Chemistry Analyzer^89^. Non-HDL-C was calculated by subtracting HDL-C from TC. Lipoprotein particle concentrations were measured by gas-phase electrophoresis (ion mobility)^90^ according to the method of Caulfield et al.^91^

### Histological analyses

Frozen liver samples were embedded in Tissue-Tek optimum cutting temperature (OCT) compound on frozen blocks and sectioned on a Cryostar NX60 Cryostat Instrument (Epredia). Slides were sectioned at 10 µm thickness, stained with Oil Red O (ORO) or H&E, and scanned using a Versa 200 Automated Slide Scanner (Leica) with a 20 x air objective lens. Images were analyzed by ImageJ.

### Statistical analysis

All data are presented as the mean ± standard error of mean (SEM). *N*-values in the figures refer to biological replicates and at least 3 replicates were conducted per condition and experiment. *P*-values were calculated using Student’s t-tests for two groups. To compare more than two groups, one-way analysis of variance (ANOVA) with Tukey’s post hoc test were used. Analyses were performed using GraphPad Prism 9 software (GraphPad Software, Inc.) *P<0.05* was considered statistically significant.

## ACKNOWLEDGEMENTS

We thank UCSF Gladstone Histology Core for their assistance in sectioning, ORO and H&E staining, and imaging of mouse liver samples. Justin Kim and Andy Su assisted with measurements using the Agilent Seahorse XFe96 Extracellular Flux Analyzer and qPCR. Dr. Marisa Medina kindly provided use of her laboratory equipment and space.

## AUTHOR CONTRIBUTIONS

Conceptualization, N.V.Y., R.M.K., and E.T.; Methodology, N.V.Y.; Formal Analysis, N.V.Y. and M.K.; Investigation, N.V.Y., J.Y.C., T.T., K.G., S.K., J.O., J.H.O., A.C., M.K., R.Z., D.J., and P.K.; Resources, R.M.K., D.J., and J.C.; Writing – Original Draft, N.V.Y., R.M.K., and E.T.; Writing – Review & Editing, N.V.Y., J.Y.C., K.G., T.T., J.H.O., S.K., J.O., A.C., M.K., R.Z., D.J., P.K., J.C., E.T., and R.M.K.; Visualization, N.V.Y.; Supervision, R.M.K. and E.T.; Project Administration, R.M.K., Funding Acquisition, R.M.K.

## SUPPLEMENTAL FIGURES

**Figure S1.**
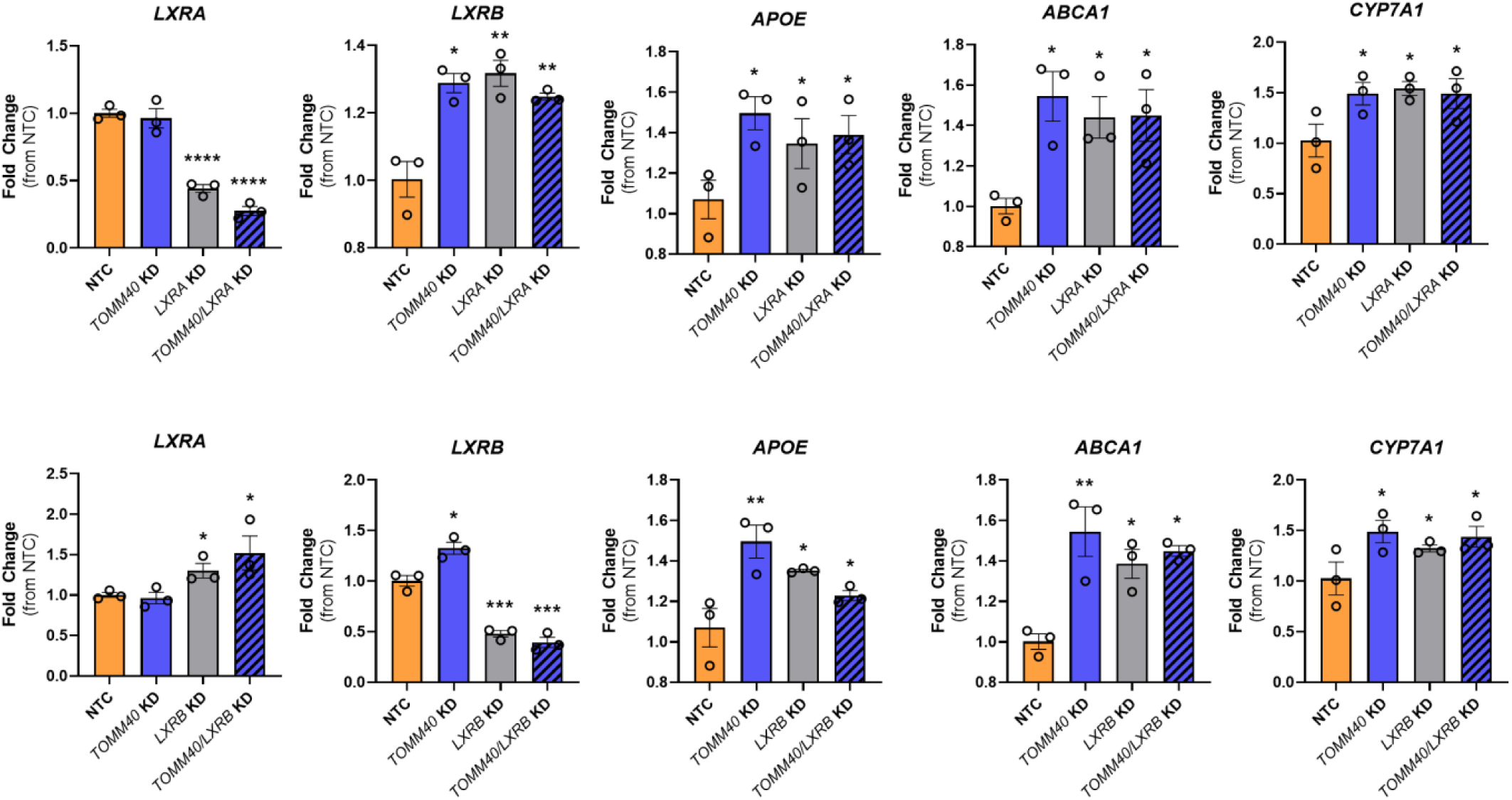
LXRA and LXRB isoforms reciprocally maintain LXR expression and regulate downstream gene targets. Relative mRNA transcript levels *of LXRA, LXRB, APOE, ABCA1,* and *CYP7A1*, compared between NTC vs. *TOMM40, LXRA, LXRB siRNA*s, singly or in combination, in HepG2 cells. *p<0.05, **p<0.01, ***p<0.005, ****p<0.001 vs. NTC by one-way ANOVA, with post-hoc Student’s t-test. (*n=3* biological replicates)

**Figure S2.**
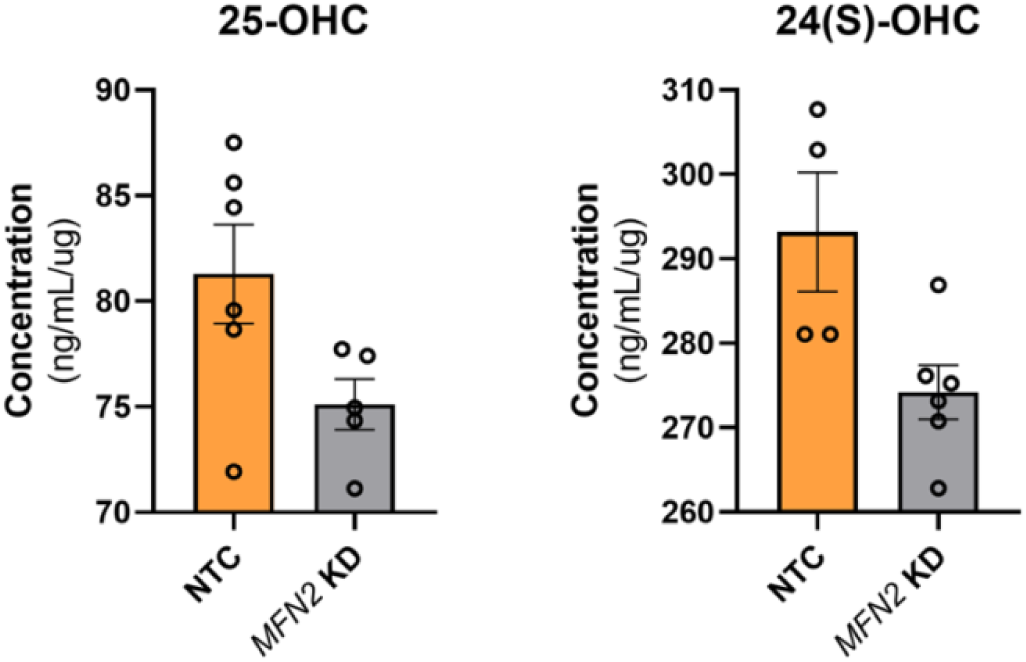
*MFN2* KD does not affect enzymatic-derived oxysterols in HepG2 cells. Analysis of enzymatic-derived 25-OHC levels and 24(S)-OHC in *TOMM40* KD vs. NTC HepG2 cells by ELISAs. (*n=3* biological replicates)

**Figure S3.**
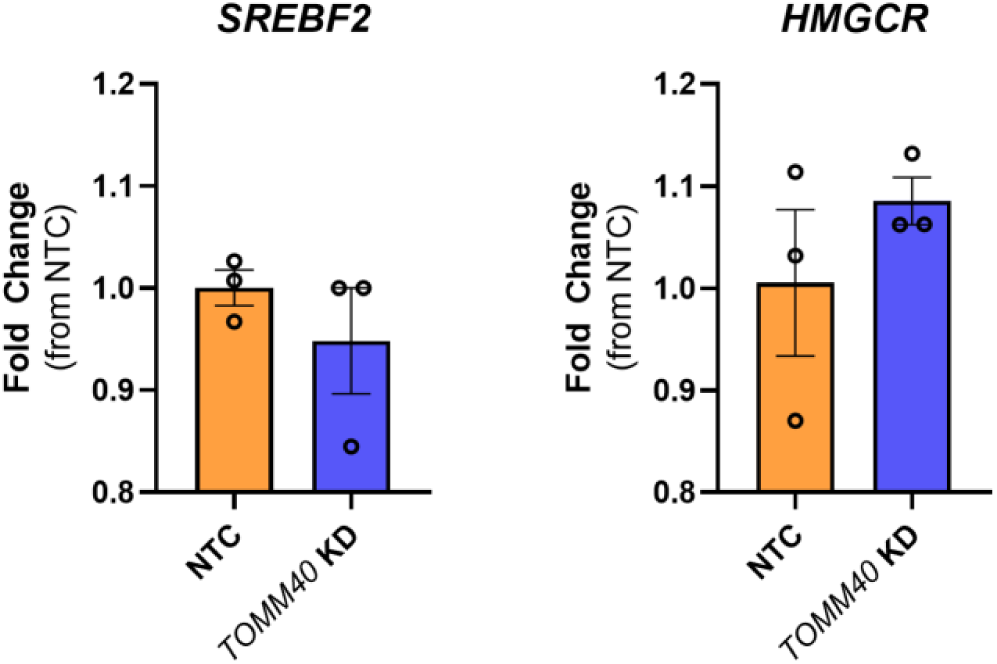
TOMM40 KD does not affects *SREBF2* or *HMGCR* mRNA transcript levels in HepG2 cells. (*n=3* biological replicates)

**Figure S4.**
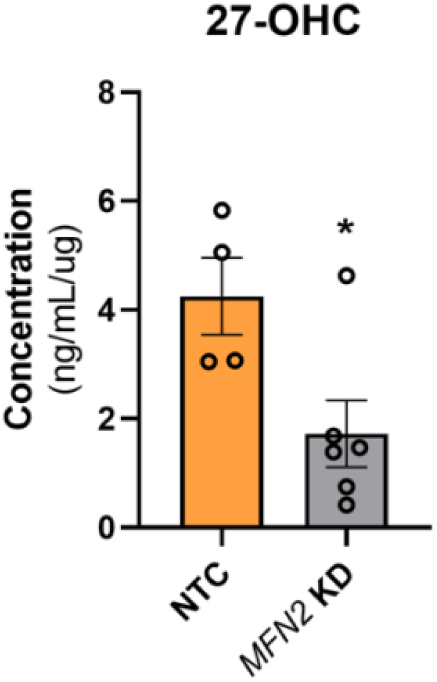
MFN2 KD reduces 27-OHC levels in HepG2 cells. Analysis of enzymatic-derived 27-OHC levels in NTC vs. *TOMM40* KD HepG2 cells by ELISA. *p<0.05 vs. NTC by one-way ANOVA, with post-hoc Student’s t-test. (n=3 biological replicates)

**Figure S5.**
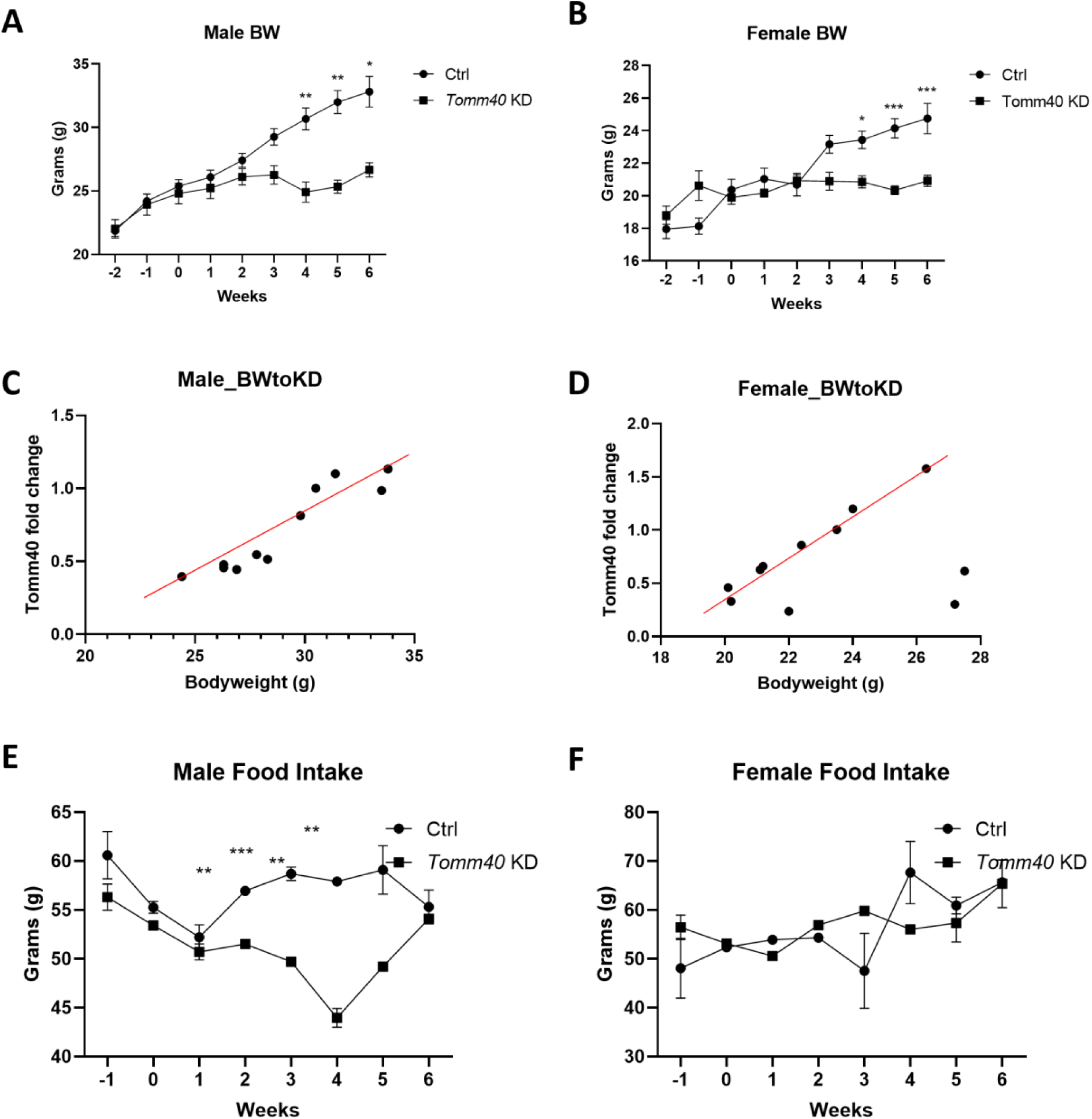
Body weight and food intake measurements of AAV8-*Tomm40* shRNA C57BL/6J mice. (A-B) Weekly bodyweight of male (A) and female (B) scrambled vs. *Tomm40* shRNA mice. (C-D) Comparison between *Tomm40* KD: bodyweight ratio in male (C) and female (D) mice. (E-F) Weekly food intake of male (E) and female (F) mice. *p<0.05, **p<0.01, ***p<0.005, ****p<0.001 vs. NTC by one-way ANOVA, with post-hoc Student’s t-test. (*n=6* mice/sex/group)

**Figure S6.**
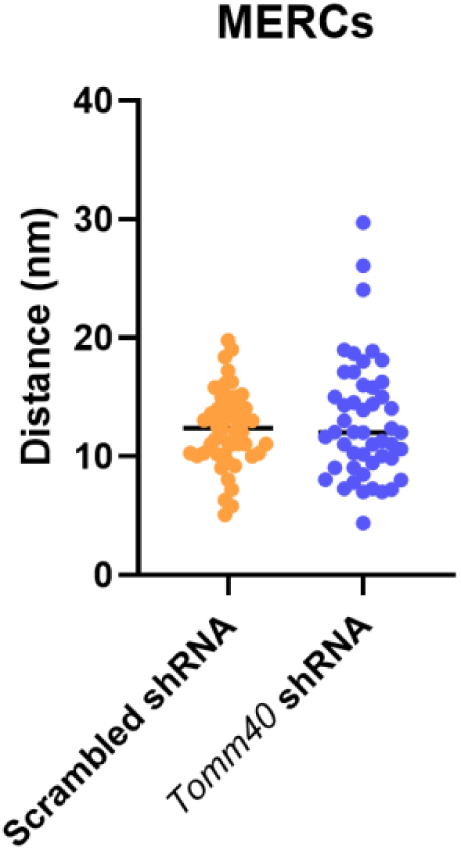
Analysis of distance (nm) between MERCs in AAV8-*Tomm40* shRNA C57BL/6J female mice liver. TEM images were analyzed by ImageJ software. (*n= 24-48* fields)

**Figure S7.**
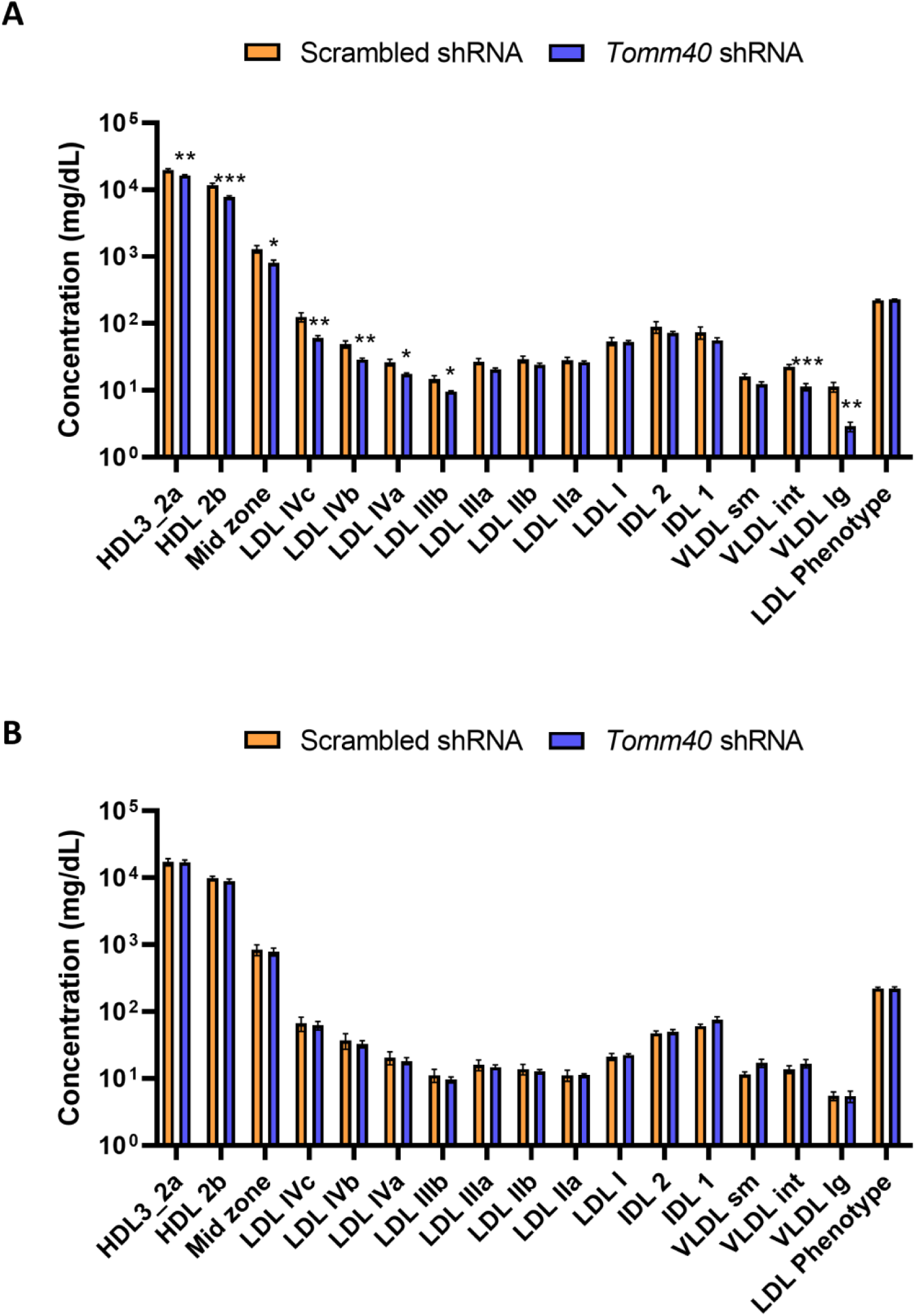
Measurements of lipoprotein particle concentrations on mouse plasma. Lipoprotein concentrations were quantified by ion mobility in (A) male and (B) female mouse plasma in *Tomm40* KD vs. scrambled control. Mouse plasma lipoprotein subfractions were analyzed based on human clinical lipoprotein particle classifications. *p<0.05, **p<0.01, ***p<0.005, ****p<0.001 vs. NTC by one-way ANOVA, with post-hoc Student’s t-test. (n=6 mice/sex/group)

**Figure S8.**
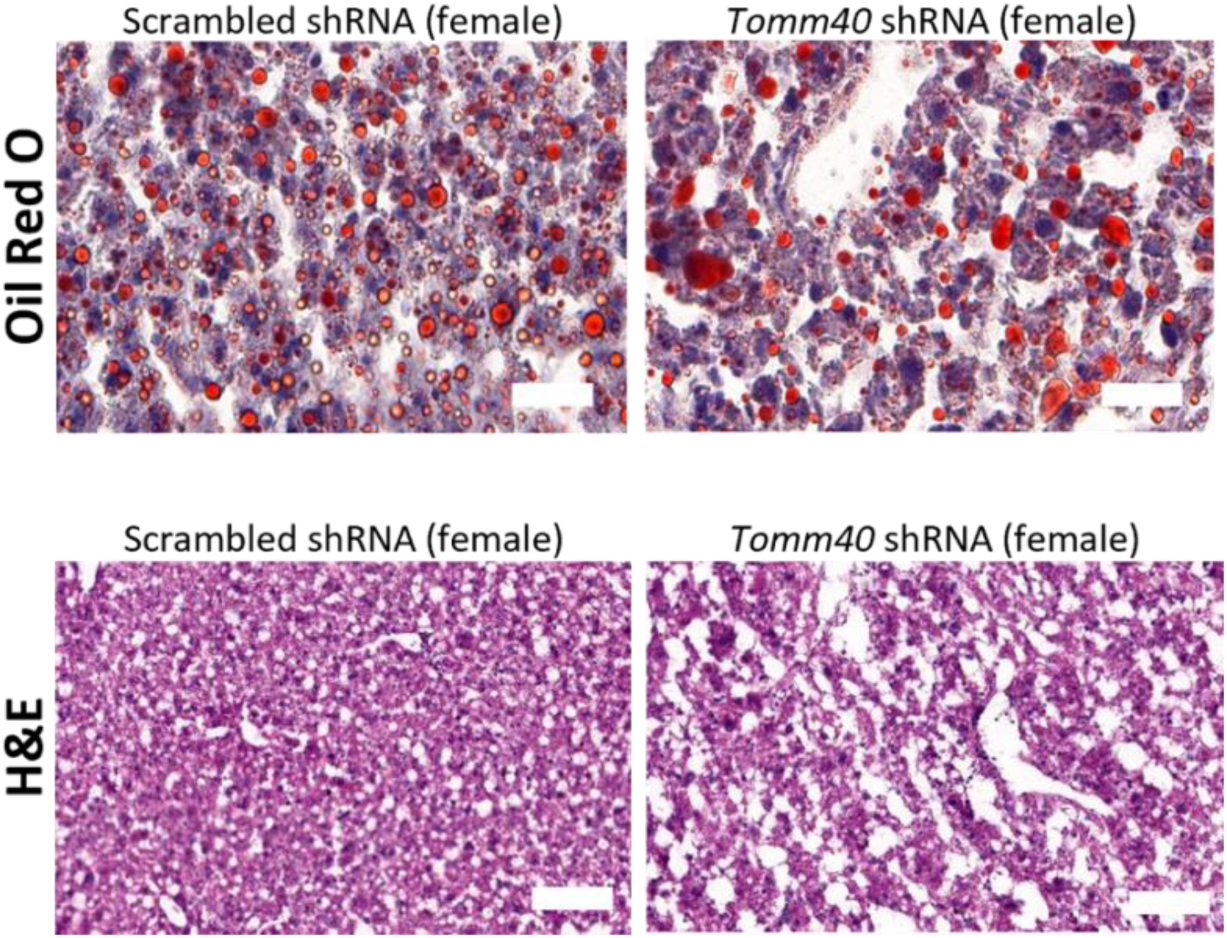
Representative Oil Red O and Hematoxylin-Eosin stained liver samples of female mice. (400 x magnification) Scale bars, 50 µm.

**Figure S9.**
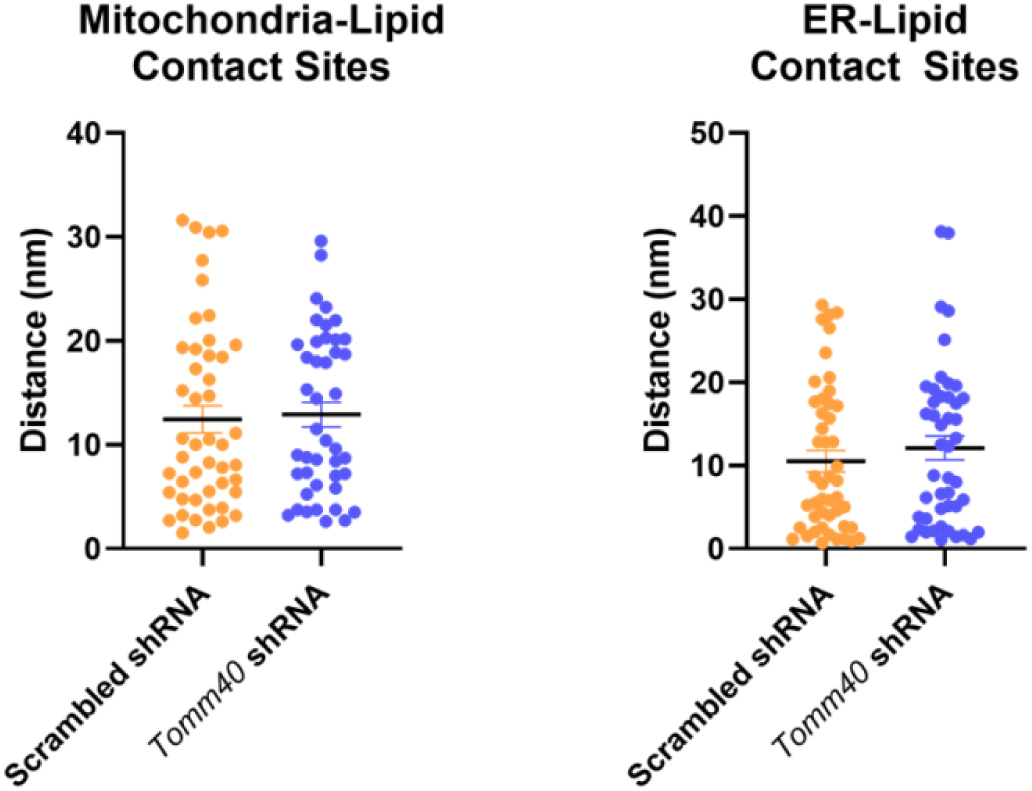
Analysis of TEM micrographs indicating no differences in lipid droplet-ER and lipid-droplet mitochondria contact sites in female mice. (A) Distance between mitochondria and lipid contact sites (nm), (B) Distance between ER and lipid contact sites (nm). (*n=24-48* fields)

**Figure S10.**
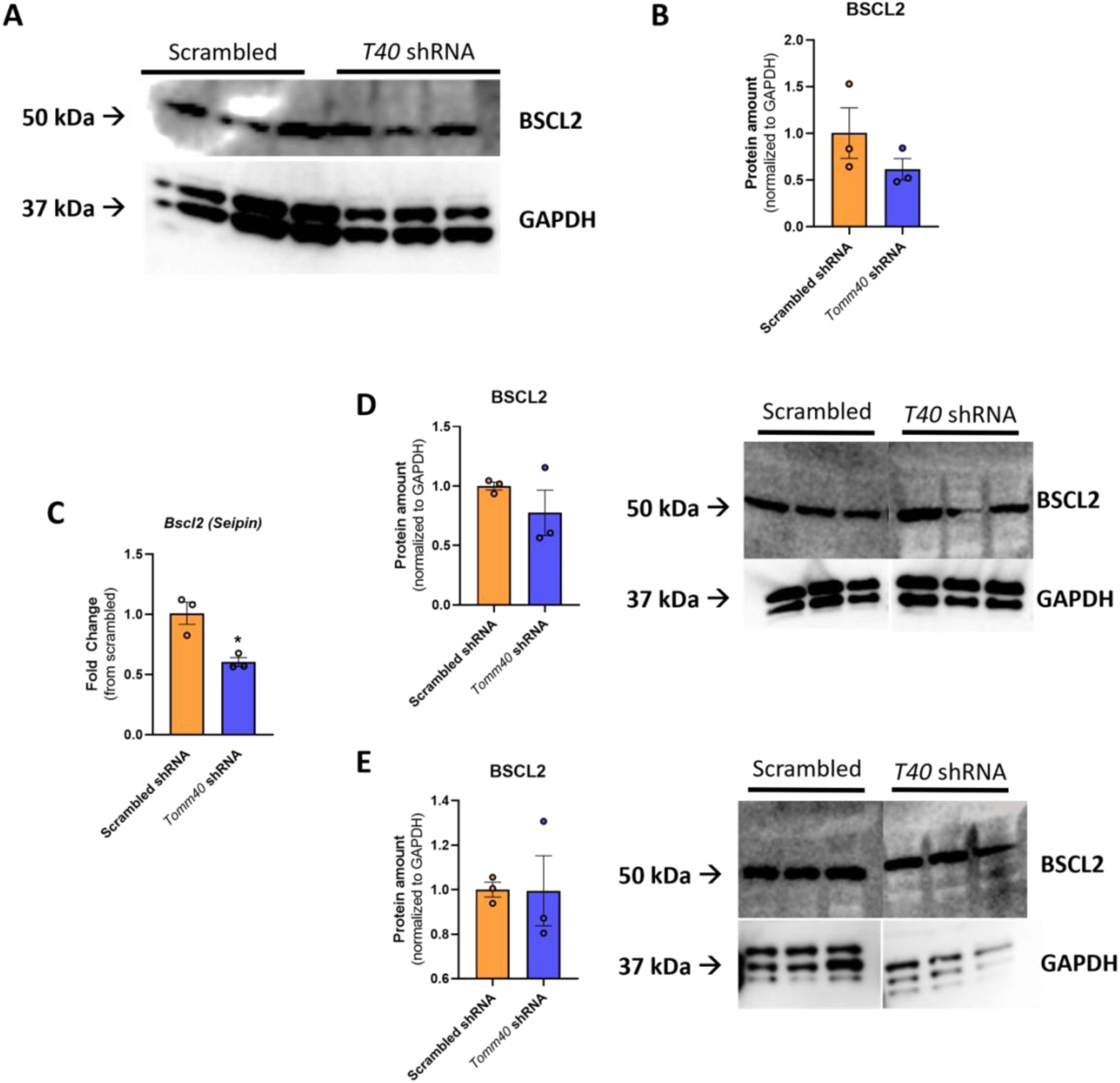
Quantification of *Bscl2*/BSCL2 expression in mouse hepatic tissues. (A) Representative western blot and (B) relative protein amount of BSCL2 protein expression in cytosolic fractions of scrambled vs. *Tomm40* shRNA male mice liver. (C) mRNA transcript levels of Bscl2 in scrambled vs. *Tomm40* shRNA female mice liver. Relative protein amount and representative western blot of BSCL2 protein expression in (D) mitochondria-associated membranes (MAMs) and (E) cytosolic fractions in scrambled vs. *Tomm40* shRNA female mice livers. For all: *n=3* mice per group; *p<0.05, vs. scrambled shRNA by one-way ANOVA, with post-hoc Student’s t-test to identify differences between groups. Data are represented as mean ± SEM.

**Figure S11.**
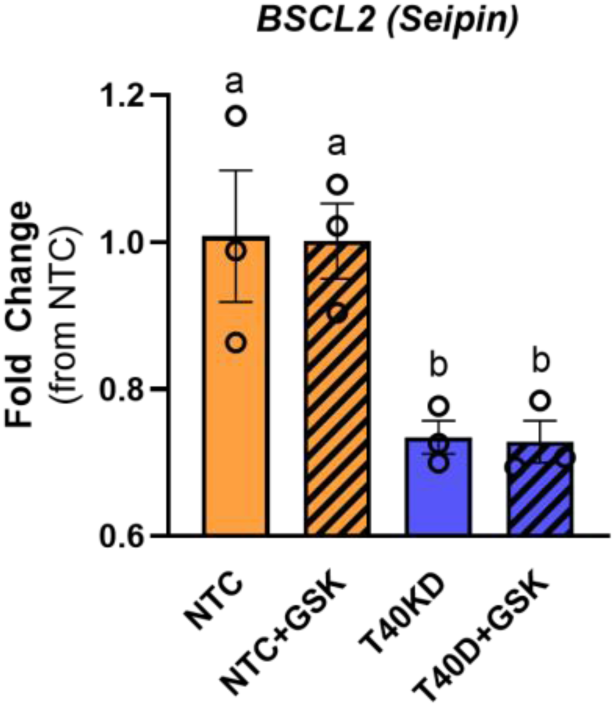
mRNA transcript levels of *BSCL2* in HepG2 cells. mRNA transcripts of *BSCL2* from NTC vs. TOMM40 KD HepG2 cells with or without addition of GSK2033 were quantified by qPCR. p<0.05 for *a* vs. *b* vs. *c* vs. *d* by two-way ANOVA, with Sidak’s multiple comparisons test. (n=3 biological replicates)

**Figure S12.**
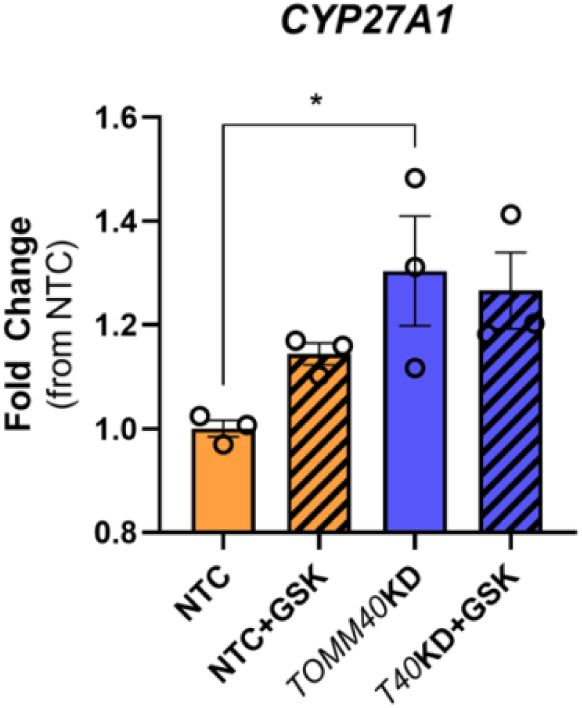
CYP27A1 mRNA transcript levels. mRNA transcripts of *CYP27A1* from NTC vs. TOMM40 KD HepG2 cells with or without addition of GSK2033 were quantified by qPCR. *p<0.05 vs. NTC by one-way ANOVA, with post-hoc Student’s t-test. (n=3 biological replicates)

